# Dynamic fibrillar assembly of αB-crystallin induced by perturbation of the conserved NT-IXI motif resolved by cryo-EM

**DOI:** 10.1101/2024.03.22.586355

**Authors:** Russell McFarland, Steve Reichow

**Affiliations:** Department of Chemical Physiology and Biochemistry, Oregon Health & Science University, Portland, Oregon 97239, USA; Vollum Institute, Oregon Health & Science University, Portland, Oregon 97239, USA; Department of Chemistry, Portland State University, Portland, Oregon 97201, USA; Department of Biochemistry & Molecular Genetics, University of Colorado School of Medicine, Aurora, Colorado 80045

**Keywords:** α-crystallin, HSPB5, CRYAB, small heat shock protein (sHSP), chaperone, proteostasis, fibril, structural biology, cryo-electron microscopy (cryo-EM), light-scattering, cataract, α-crystallin-opathies

## Abstract

αB-crystallin is an archetypical member of the small heat-shock proteins (sHSPs) vital for cellular proteostasis and mitigating protein misfolding diseases. Gaining insights into the principles defining their molecular organization and chaperone function have been hindered by intrinsic dynamic properties and limited high-resolution structural analysis. To disentangle the mechanistic underpinnings of these dynamical properties, we mutated a conserved IXI-motif located within the N-terminal (NT) domain of human αB-crystallin. This resulted in a profound structural transformation, from highly polydispersed caged-like native assemblies into a comparatively well-ordered helical fibril state amenable to high-resolution cryo-EM analysis. The reversible nature of the induced fibrils facilitated interrogation of functional effects due to perturbation of the NT-IXI motif in both the native-like oligomer and fibril states. Together, our investigations unveiled several features thought to be key mechanistic attributes to sHSPs and point to a critical significance of the NT-IXI motif in αB-crystallin assembly, dynamics and chaperone activity.

## INTRODUCTION

Small heat-shock proteins (sHSPs) are a conserved family of protein chaperones, critical for maintaining cellular proteostasis^1–3^. These holdases recognize and sequester destabilized proteins (*aka* clients), preventing detrimental aggregation events in an ATP-independent manner. The sequestered client remains in a refolding-competent state that may be rescued by ATP-dependent chaperones, like the HSP70 system^4^. Ten sHSP proteins (HSPB1-10) are found in human that are differentially expressed throughout the body^5,6^. HSPB5 (aka, αB-crystallin; CRYAB) is considered an archetype of the mammalian sHSP family and is ubiquitously expressed, with high levels in the eye lens, cardiac and neuronal tissues^7,8^. Because of its critical physiological roles, aberrant function or dysregulation of αB-crystallin is associated with a variety of protein misfolding diseases, including cataract, Alzheimer’s disease, Parkinson’s disease, neuromuscular disease, as well as some cancers^9,10^.

Despite the physiological significance and involvement in various diseases linked to sHSPs, our grasp of these processes remains constrained by the insufficiency of high-resolution structural insights into this system^11,12^. This limitation stems from the complex dynamics exhibited by eukaryotic sHSPs that can form high-order assemblies. Under normal conditions, αB-crystallin forms a continuum of large polydispersed oligomers (∼10–40 subunits)^13–16^, characterized by rapid subunit exchange^17–20^. These inherent dynamics are pivotal to the chaperone mechanism, enabling sHSP’s to effectively scavenge and sequester a diverse range of client proteins^21^. At the subunit level, αB-crystallin (20.2 kDa) features a tripartite domain organization seen in other sHSPs^22,23^. Central to this is a conserved α-crystallin domain (ACD, ∼80 residues in αB-crystallin) shared by all sHSPs. This domain acts as a structural hub, flanked by a variable N-terminal domain (NTD, ∼65 residues in αB-crystallin) and C-terminal domain (CTD, ∼28 residues in αB-crystallin) that are both highly dynamic and flexible^24–26^.

The NTD and CTD endow many sHSPs with the capacity to form polydispersed high-order oligomeric structures. Truncated mutants that lack these domains form stable ACD dimers, considered the fundamental building blocks of sHSPs that have been well-characterized structurally^27–31^. Within αB-crystallin and other eukaryotic sHSPs, the CTD performs a well-established role in oligomerization, a process facilitated by a conserved ‘IXI-motif’ (with X indicating a variable sequence position), followed by an extension of polar and charged residues that enhance overall solubility^32–34^. The influence of the CT-IXI motif on oligomer assembly involves domain-swapping interactions, including binding to a hydrophobic groove located in the ACD.

While the NTD shows considerable sequence variability across sHSPs, it contains regions that remain relatively conserved among eukaryotic sHSPs that exhibit a high degree of hydrophobicity and similarly involved in multiple interactions with the ACD within the context of native oligomeric formations^24,35,36^. The functional role of the NTD is less clearly defined but has been acknowledged for its involvement in client binding and specificity, as well as oligomer assembly^21,37,38^.

Notably, the NTD in αB-crystallin contains an additional IXI-motif located near the beginning of the polypeptide chain (residues 3-5). This N-terminal motif is also present in other sHSPs, such as HSPB3, HSPB4/αA-crystallin, and HSPB6 (**Fig. 1a**), where isoleucine is sometimes found as valine. Current models suggest the NT-IXI competes with the CT-IXI motif for binding to the hydrophobic groove at the ACD, contributing to subunit exchange dynamics and oligomer assembly^36^. Due to these various forms of dynamics, structural investigations of full-length αB-crystallin and other high-order sHSPs have so far been constrained to relatively low-resolution or integrative models^39–42^.

**Figure 1.**
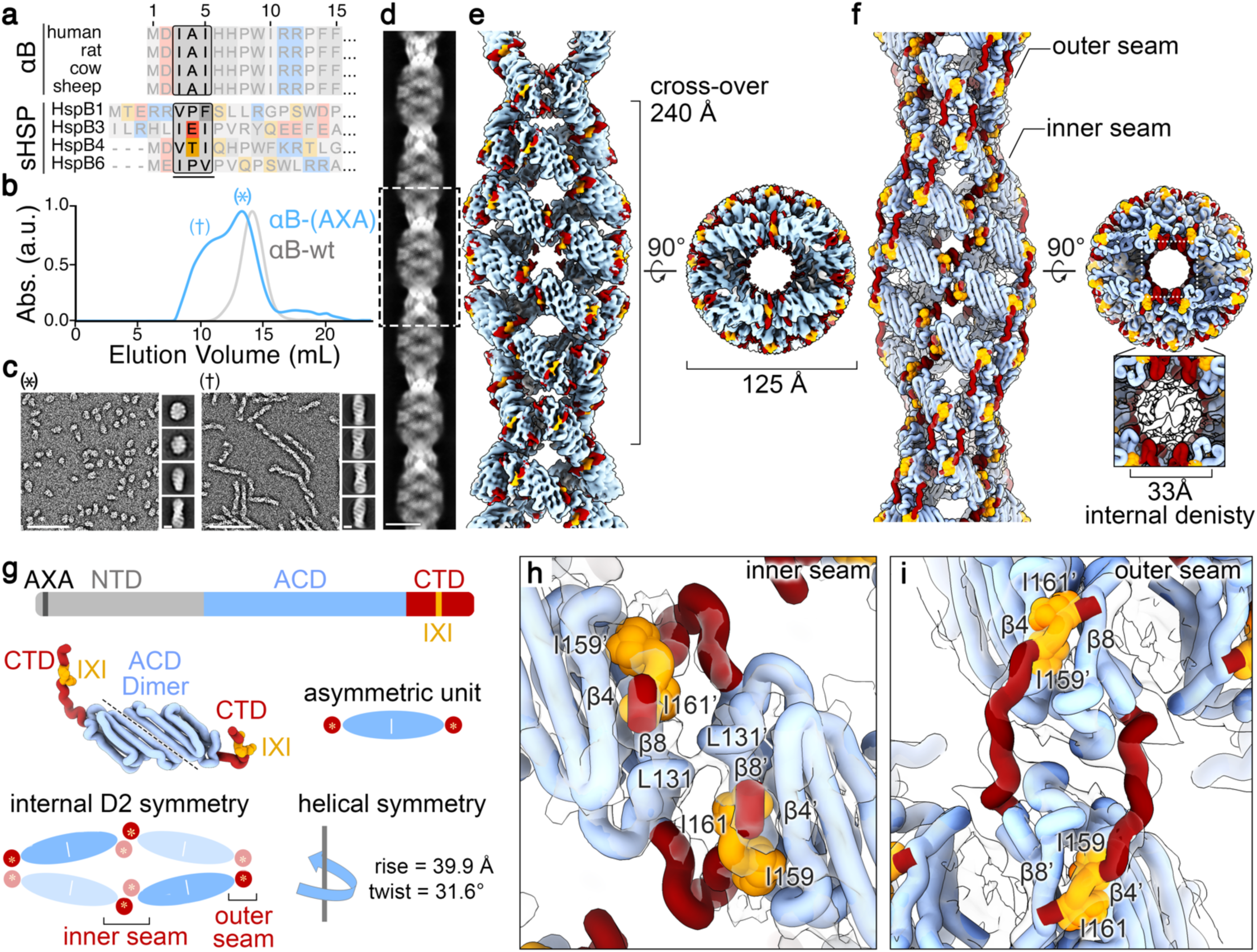
αB-AXA fibril resolved by cryo-EM. **a.** Sequence alignment of the distal region of the N-terminal domain of αB-crystallin showing conservation of the NT-IXI motif (boxed) across model species that was mutated in this work (top), and putatively identified NT-IXI regions in other related sHSPs (bottom). In canonical IXI-motifs, isoleucine may be replaced by valine. In other sHSPs, such as HSPB1, the NT-IXI motif may be more cryptic, with isoleucine positions replaced by other hydrophobic residues. **b**. Size-exclusion chromatography (SEC) profile of αB-wt (grey trace) and αB-AXA (blue trace). Peak fractions are labeled as (†) indicating region containing fibrils and (*) with native-like assemblies. **c.** Representative EM micrographs (scale = 100 nm) and 2D classes of αB-AXA samples (scale = 10 nm) isolated by SEC fractions (*, left) and (†, right). **d.** Montage of 2D classes obtained by cryo-EM (scale = 10 nm). **e.** Cryo-EM density map of the αB-AXA fibril state resolved at 3.4 Å resolution and **f.** atomic model displayed in cartoon representation, with the ACD in blue, CTD in red and CT-IXI in orange. **g.** Illustration of domain organization, colored as in panel f, with NTD colored in grey (top), structure and illustration of the asymmetric unit showing 2-fold internal pseudosymmetry of the ACD dimer (middle), and schematic illustrating the internal D2 and helical symmetries present in the fibril assembly (bottom). **h, I.** Zoomed views of CT-IXI motif binding with the ACD β4/8 groove at inner and outer seam, respectively. Cryo-EM density is displayed in transparency. Interacting residues between neighboring subunits are labeled.

In this study, we aimed to explore the impact of disrupting the competition between the CT-IXI and NT-IXI motifs through mutational ablation of the NT motif. Our hypothesis was that silencing the NT-IXI motif might reduce polydispersity, making the system more amenable to high-resolution structural analysis. Surprisingly, introducing an NT-AXA variant into human αB-crystallin (αB-AXA), where conserved isoleucines were replaced by alanines, led to the emergence of ordered helical fibril assemblies formed during cellular expression. The isolated fibrils are non-amyloid in nature and found to be fully reversible with native-like caged oligomers upon mild temperature changes (25–42° C), with fibrils favored at lower temperatures. Cryo-electron microscopy (cryo-EM) provided high-resolution structural and mechanistic insights, resolving the fibril assembly at ∼3.4 Å resolution, enabling detailed atomic modeling. The fibril state of αB-crystallin was found to retain essential features of native sHSP oligomers, including roles of the NTD and CTD in high-order assembly, rapid subunit exchange dynamics, and significant conformational plasticity. The reversible nature of the αB-AXA variant allowed for a functional dissection of chaperone activity in both the fibril and native-like oligomer states, unveiling varying degrees of impairment in chaperone activity compared to wildtype when tested against the model aggregating client, lysozyme. These findings offer long-sought insights into the fundamental principles governing sHSP assembly and dynamics, shedding light on the pivotal role of the conserved NT-IXI motif in αB-crystallin and its dynamic interplay with the CT-IXI.

## RESULTS

### Disrupting the conserved NT-IXI motif transforms αB-crystallin into helical fibrils

The NT-IXI motif in αB-crystallin, represented as IAI, is well conserved among model mammalian species (**Fig. 1a** and **Extended Data Fig. 1**). To comprehensively assess the structural and functional implications of this duplicate IXI-motif, we engineered a variant of the human αB-crystallin gene, introducing alanine substitutions in place of the conserved isoleucines. This NT-AXA variant (αB-AXA) was expressed in bacterial cells and purified without the use of purification tags or other changes to the gene sequence, using a procedure adapted for wildtype αB-crystallin (αB-wt) (**Fig. 1b** and **Extended Data Fig. 2**)^1,16^. In consideration of the sensitivity of αB-wt oligomerization to environmental conditions, including factors like pH and divalent cations^43^, the ensuing structural and functional investigations were conducted in buffer conditions containing 20 mM HEPES (pH 7.4), 150 mM NaCl, and 1 mM EDTA.

Size-exclusion chromatography (SEC) analysis of isolated αB-AXA initially revealed a significant degree of enhanced polydispersity and potential aggregation. The elution profile of αB-AXA exhibited a major peak that coincided with the elution profile of αB-wt (* in **Fig. 1b**). Additionally, an overlapping broad peak was observed, extending towards the high molecular weight void volume of the SEC column († in **Fig. 1b**). Further examination by electron microscopy (EM) of negatively stained samples extracted from these primary elution peaks identified two predominant species. The fraction showing elution overlap with αB-wt mainly consisted of spherical oligomers, measuring approximately 14–16 nm in diameter (**Fig. 1c**, *), consistent with previous characterizations of αB-wt^16,41,44,45^ (see also **Extended Data Fig. 2**).

Interestingly, the high-molecular weight fractions exclusively displayed fibrillar structures with distinct helical features (**Fig. 1c**, †). Employing reference-free two-dimensional (2D) class-averaging techniques, the helical fibril dimensions were determined to have a diameter of about 15 nm and a helical periodicity of around 45 nm. Striated features could be clearly resolved, arranged transversely to the principal helical axis of the fibrils, with an interval of approximately 5 nm. Within the high-molecular weight SEC fractions, fibrils exhibited varying lengths, extending up to ∼250 nm. Notably, 2D class-averaging analysis applied to the lower-molecular weight fractions uncovered the presence of short fibril assemblies (or protofibrils) in addition to the native-like oligomeric species (comparison of top and bottom 2D classes in **Fig. 1c**,*).

### High-resolution structural analysis of αB-crystallin NT-AXA fibrils by cryo-EM

The distinctive order observed in the αB-AXA fibrils suggested the potential for subjecting them to high-resolution structural analysis by cryo-electron microscopy (cryo-EM). To achieve this, we conducted further biochemical optimization of αB-AXA fibrils, as described in detail below and in the Methods section. The acquired datasets underwent image processing routines utilizing a combination of helical and single-particle methodologies implemented in cryoSPARC^46^, resulting in a final refined density map resolved to a global resolution of 3.4 Å, and local resolutions ranging from 2.5–3.5 Å (**Fig. 1d,e**; **Extended Data Fig. 3 and 4;** and **Extended Data Table 1**)

The dimensions of the resolved fibrils have a diameter of approximately 125 Å, with a refined helical periodicity measuring 480 Å, characterized by a rise of 39.9 Å and twist of 31.6° for the repeating asymmetric unit. The overall quality of the map facilitated *de novo* construction of a model covering the entire ACD (*blue, residues 75-148*) and the majority of the CTD (*red and orange, residues 149-161*) (**Fig. 1f,g**; **Extended Data Fig. 1 and 4; Supplemental Movie 1**). The last 14 residues could not be fully resolved, presumably due to the high-degree of flexibility at the c-terminus and were not modeled.

The fundamental building block of the αB-AXA fibril assembly is the ACD dimer, where each subunit adopts an expected IgG-like β-sandwich fold. In this configuration, formation of the symmetric ACD dimer occurs through an in-register interface involving β6/7 (*aka* the AP_II_ state)^27,29,39^. The conformation of the ACD dimer closely resembles previously described structures obtained from diverse αB-crystallin constructs, particularly resembling the original ACD X-ray structure (PDB 2WJ7)^27^, with an overall Cα r.m.s.d. of ∼1.6 Å (**Extended Data Fig. 5**).

Further investigation revealed several other shared features to previously described αB-crystallin ACDs. The loop that links β5 and β6 is resolved adopting a so-called upward conformation^31^. This conformation is supported by electrostatic interactions between D109 and R116 as well as R120 from the adjacent protomer, effectively reinforcing the dimer interface (**Extended Data Fig. 5**). Moreover, the β2 strand is resolved for only one of the protomers in the ACD dimer (subunit A, forming the outer seam), reflecting differences in the overall stability of this secondary structural element (**Extended Data Fig. 1 and 5**). This β-strand is part of the ‘boundary region’ and has not consistently been resolved in various X-ray and NMR structures of the ACD^27,36,39^. Here, the differences in conformational stability in this region could stem from the pseudo-symmetric relationship between the two protomers and possibly coupled with variations of interaction with NTDs, as further elaborated below.

In the context of the helical assembly, each ACD dimer is arranged with overall D2-symmetry (**Fig. 1g**), that establishes a two-start helical arrangement and leads to the rung-like features created by corresponding pairs of ACD dimers. The helical architecture generates an interior cavity with approximate diameter of 5 nm that is filled with poorly defined density, presumably belonging to the NTD. The helical rise of 39.9 Å generates separation between the ACD rungs, resulting in the formation of ∼20 Å wide fenestrations that are clearly resolved along the helical assembly (**Fig. 1e,f**). Notably, these features bear resemblance to the fenestrated cage-like architectures previously proposed from low-resolution cryo-EM models of wildtype αB-and αA-crystallins, and other sHSPs^40,42,47–50^.

### The CT-IXI forms multiple domain-swapping interactions with the ACD groove

The internal symmetry of the fibril assembly gives rise to two distinct interfaces formed between the ACD dimer building blocks, referred to as the inner and outer seam (**Fig. 1f-i**). The inner seam emerges from the pairing of ACD dimers that constitute the rungs of the assembly, which appear stitched together through domain-swapped CTDs exchanged between adjacent subunits (**Fig. 1g,h**). Conversely, the outer seam is formed along the edge of the filament, where ACD dimers are stitched together along the helical axis through domain swapped CTD interactions occurring between stacked subunits (**Fig. 1g,i**). As a result, this structural arrangement gives rise to two distinct conformational states adopted by the CTD.

In both scenarios, the domain swap interactions involve participation of the conserved CT-IXI motif from one protomer and the hydrophobic groove generated by β4/8 from a protomer on a neighboring ACD. In the context of the inner seam, the CT-IXI is positioned such that I159 establishes interactions with residues V91, V93, and I133, while I161 interacts with L89, V91, L137, and L143 that form hydrophobic pockets within β4/8 groove (**Fig. 1h** and **Extended Data Fig. 6**). S135, situated within the ACD groove, lies between these two hydrophobic pockets, effectively bifurcating the two bound isoleucine residues. This knob-and-hole interaction mechanism is analogous to previously structural studies of the αB-crystallin ACD bound to CTD-mimicking peptides^30,34^. Remarkably, the only other notable interaction stabilizing this interface occurs between symmetry-related residues L131, contributed by adjacent ACD domains (**Fig. 1h**).

In comparison, the CT-IXI interaction for the outer seam occurs in the opposite orientation with respect to the ACD. In this case, I159 inserts into the pocket formed by L89, V91, L137, and L143, while I161 engages with the pocket formed by V91, V93, and I133 (**Fig. 1i** and **Extended Data Fig. 6**). The flexibility of the linker connecting the ACD to the CT-IXI, coupled with the palindromic sequence within this region of the CTD, allows for a bidirectional orientation about the ACD. The CT-linker adopts a distinctive compact turn to facilitate the angle of approach for the inner-seam binding site. Conversely, the outer-seam interaction relies on a relatively extended conformational state for the CT-linker, positioning the IXI-motif against the interacting ACD β4/8 groove with a reversed orientation compared to the inner-seam interaction.

### A quasi-ordered NTD buttresses the interior core of the ACD fibril assembly

The NTD is considerably less resolved in the cryo-EM map, as compared to the ACD and CTD domains, and appears as a quasi-ordered density that occupies nearly the entire interior cavity of the fibril (**Fig. 2a**). The poor resolvability of this domain may reflect inherent disorder or the assumption of multiple quasi-ordered conformational states. These states, despite substantial efforts employing various image processing techniques, could not be fully resolved or differentiated by image classification routines.

**Figure 2.**
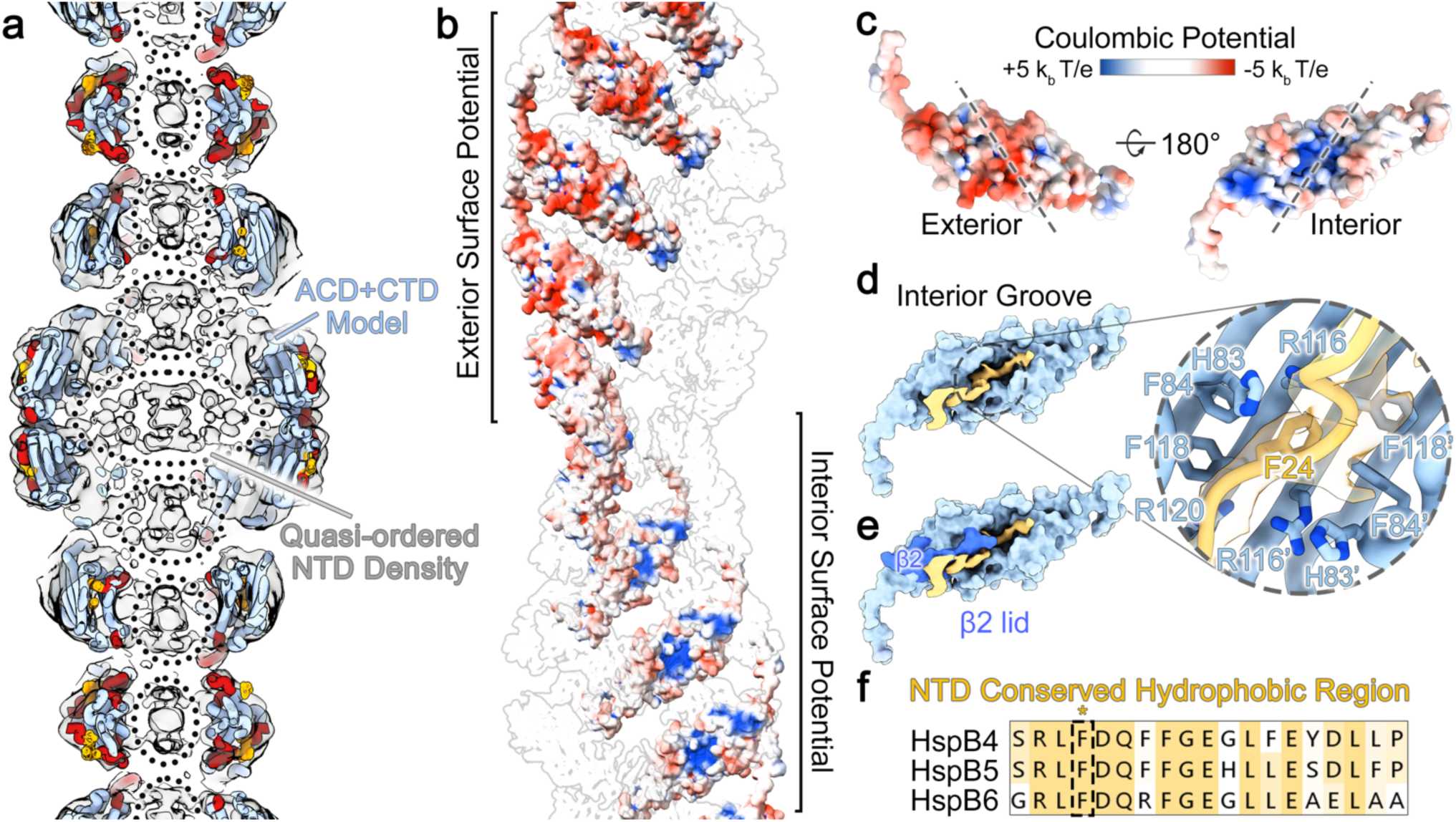
The quasi-ordered NTD fills an internal cavity of the αB-AXA fibril and forms interactions with the ACD. **a.** Cut-away view showing αB-AXA fibril model colored as in Fig. 1, with the cryo-EM density map overlayed in grey. The unmodeled density filling the interior cavity of the fibril is assigned to the NTD (dotted circles). **b.** Electrostatic surface representation of αB-AXA fibril, with only a single filament stand displayed for clarity (red: −5 kb T/e, white: 0 kb T/e, blue: +5 kb T/e). **c.** Electrostatic surface representation of a single αB-AXA subunit, oriented with exterior surface (left) and interior surface (right). The ACD dimer interface is indicated by dotted lines. The β2 strand is omitted for clarity. **d.** Surface representation of the ACD dimer (light blue) and segmented cryoEM map showing a region of the NTD (yellow) bound within the interior groove of the ACD. This density is putatively assigned to a conserved hydrophobic-rich region of the NTD (residues 20-32). Zoom view shows a Cα trace fit into the putative NTD density with a conserved F24 residue displayed. Residues from the ACD that interact with this peptide region also displayed and labeled. **e.** Same as panel d, with surface representation of the β2 strand resolved in a single protomer (dark blue) that forms a lid covering the bound NTD region. **f.** Sequence alignment of the putative NTD peptide binding region, showing conservation of hydrophobic-rich region with phenylalanine (F24) boxed.

The high-order organization of the ACD dimers gives rise to two distinct electrostatic surface properties within the fibril assembly (**Fig. 2b**). The exterior surface, comprising β4, β5, and β6/7 of the ACD dimer, is relatively smooth and features an overall electronegative surface potential (**Fig. 2b,c**). In comparison, the interior surface of the helical assembly, composed of β3, β8, and β9 of the ACD dimer, displays a mix of hydrophobic regions along with a distinctive positively charged groove formed by the ACD dimer interface (**Fig. 2b,c**). This internal ACD groove is occupied by a moderately well-resolved portion of the cryo-EM map attributed to a segment of the NTD (**Fig. 2d**, yellow). The interaction appears to be reinforced by the β2 strand from subunit A of the ACD, which lays across the top of the NTD region forming a lid-like feature that may further stabilize the ACD-NTD interaction (**Fig. 2e**, dark blue).

While the local resolution of the NTD segment bound by the ACD site was insufficient for unequivocal sequence assignment, it is recognized that a conserved hydrophobic region of the NTD has been implicated in binding to the ACD interior groove region in α-crystallins and other sHSPs (**Fig. 2f** and **Extended Data Fig. 6**)^36,51,52^. We fitted this NTD region into the cryo-EM density, corresponding to αB-crystallin residues LFDQFFGE (residues 23-30) (**Fig. 2d**, *zoom*). This peptide region contains several bulky hydrophobic sidechains (F24, F27, F28) that align with the cryo-EM map features and would closely interact with other hydrophobic residues constituting the ACD groove (*e.g.,* F84, F118, and H83). However, due to the uncertainties associated with limited resolution, this sequence assignment remains tentative and was fit as poly-alanine in our final model.

This NTD peptide binding site is formed by the ACD dimer interface, which has been previously resolved by X-ray crystallography in varying registers (termed AP_I_ through AP_III_)^9^. In the αB-AXA fibril assembly, only the symmetric AP_II_ state is discernable. Since this feature falls within the asymmetric unit of the helical assembly, it is unaffected by the symmetry operations applied during image processing. Notably, alternative registers would be incompatible with the observed NTD peptide binding interactions proposed above, and may therefore reinforce this specific ACD register. In a broader context, these observations implicate a critical role of the NTD in buttressing the overall architecture of the ACD core assembly, which is otherwise supported by domain swapped CTD interactions that are interconnected solely through flexible linkers.

### αB-AXA fibril state retains a high-degree of conformational plasticity

Initial attempts at refinement of the αB-AXA fibril structure yielded 3D reconstructions with limited resolutions, in the range of ∼6–8 Å (**Extended Data Fig. 3**). It was suspected that this limited resolution might be attributed to the presence of long-range conformational heterogeneity. This notion was supported by observation that fibrils exhibit a substantial degree of flexibility, as evident from low-magnification images (**Extended Data Fig. 2**). To quantitatively evaluate this morphological flexibility, image segments were extracted from the cryo-EM dataset using large box sizes, sufficient for sampling at least two cross-over points (or one complete helical period) and subjected to 2D classification analysis (**Fig. 3a,c**). Measurement of cross-over distances resolved in the 2D classes revealed significant variation, falling within the range of approximately 230–245 Å or a variance of approximately 30 Å per helical period. At a short-range level, this distribution indicates a variation in rung-to-rung distances of only ∼1.0–1.5 Å. Likewise, distinct variability along the principal helical axis is evident in the 2D class averages, manifested with bend angles ranging from 0° to approximately ±20°, as measured over a single helical period. This conformational heterogeneity was also identified and visualized using 3D Variability Analysis (3DVA) in cryoSPARC^53^, which illuminated two primary components of structural variability characterized as a combination of stretching and bending modes (**Fig. 3b,d; Extended Data Fig. 7; and Supplemental Movie 2**).These analyses underscore the significant conformational plasticity that is retained by the helical assembly, which is congruent with the flexibility of the linkers connecting the ACD and CTD, as well as the quasi-ordered nature of the NTD that reinforces the interior of the fibril architecture.

**Figure 3.**
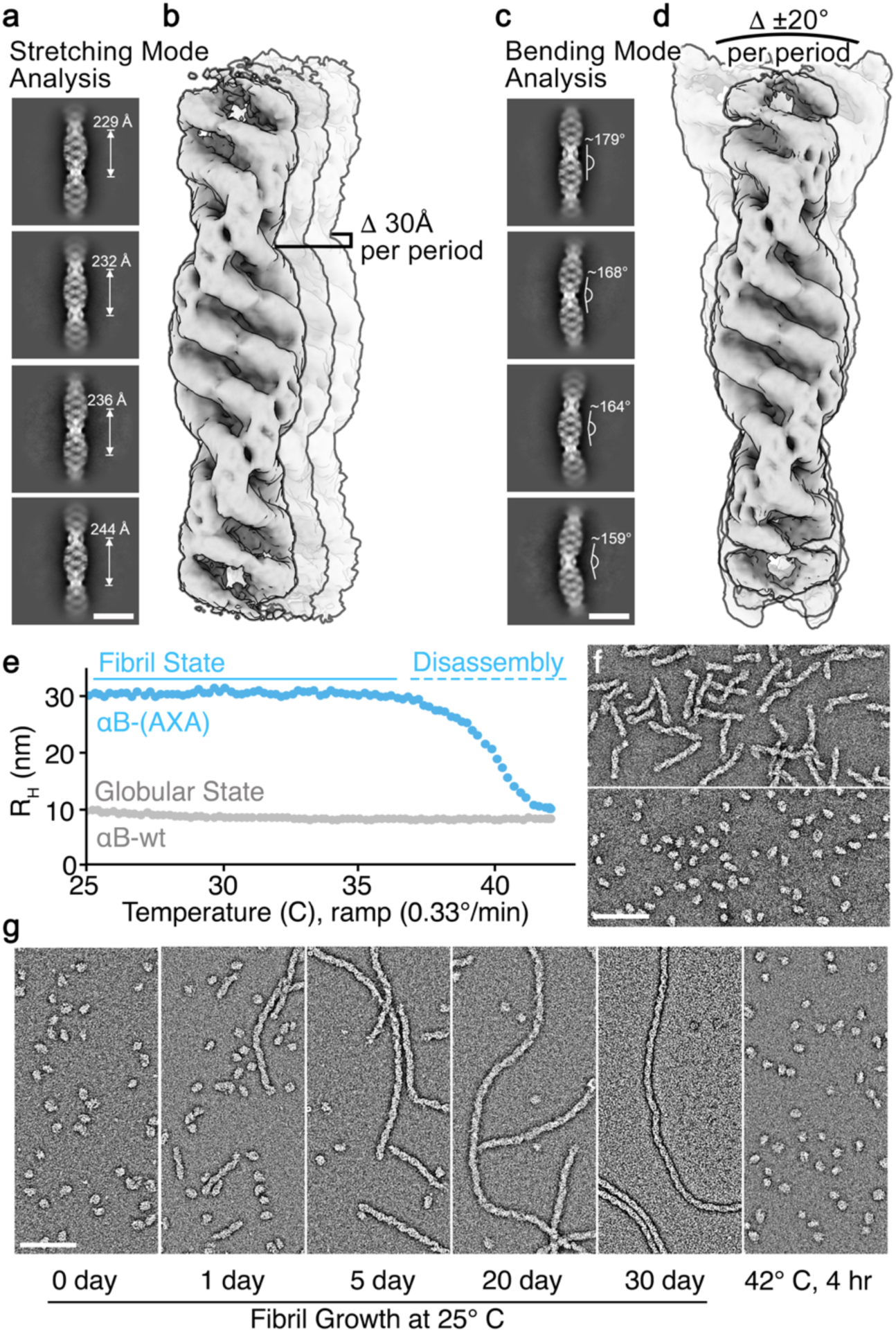
αB-AXA fibrils display structural plasticity and temperature-dependent reversibility with the native-like oligomeric state. **a.** Representative 2D class averages of αB-AXA fibrils obtained by cryo-EM showing variability in cross-over distances with measurements labeled (scale = 20 nm). **b.** Overlay of cryo-EM maps obtained from one of the primary principal components obtained by 3D variability analysis in cryoSPARC^53^, illustrating a stretching mode. A range of ∼30 Å indicates the variability of fibril length per helical period, as measured in 2D class averages. **c.** Representative 2D class averages showing variability in bend angles with measurements labeled (scale = 20 nm). **d.** Overlay of cryo-EM maps obtained from one of the primary principal components obtained by 3D variability analysis, illustrating a bending mode. A range of bend angles of ∼20° per helical period is indicated, as measured from the 2D class averages. Cryo-EM maps in panels b and d were aligned to the bottom rung density of the fibril for visualization purposes. **e.** Reversibility of αB-AXA fibrils demonstrated by conversion to the native-like globular state by monitoring the change in hydrodynamic radius (R_H_) measured by dynamic light scattering as a function of temperature. αB-wt did not show appreciable changes in R_H_ (gray trace) over temperature range of 25 – 45° C. **f.** EM micrographs of negatively stained specimens showing the starting αB-AXA fibrils (top) and resulting conversion to native-like oligomeric assemblies following heat ramp to 45° C (bottom). **g.** Native-like oligomers (left) were then incubated at room temperature (25° C) and shown to slowly convert back to the helical fiber state over a period of ∼30 days (middle panels), as monitored by EM. Grown fibrils were then readily converted back to native-like oligomers upon incubation at 42° C for four hours (right). Scale bars = 100 nm, in panels b, c.

### αB-AXA fibrils are temperature dependent and reversible with native-like caged oligomers

As described above, during the purification of αB-AXA from bacterial expression, a mixture of native-like oligomers and helical fibril states was obtained. Upon further investigation, it was noted that when the purified mixture was incubated at room temperature, a gradual transformation into the fibril state occurred. We aimed to comprehensively characterize this phenomenon by closely monitoring the transition between native-like oligomers and helical fibril morphologies, using dynamic light scattering (DLS) and by direct visualization using EM at varying incubation temperatures.

To exemplify this characterization, we considered the case of starting with a sample exclusively composed of fibrils, obtained via SEC separation from the initial mixture. These fibril structures remained stable for several days at room temperature in a standard buffer at pH 7.4, containing 150 mM NaCl and 1 mM EDTA (the same buffer utilized for structural analysis). However, when subjected to a temperature ramp ranging from 25° to 45° C over a span of 60 minutes (∼0.33°C per minute), the sample readily converted into a relatively uniform population of globular native-like assemblies (**Fig. 3e,f**). Under these conditions fibril disassembly occurs rapidly (< 10 min), with complete conversion occurring at a temperature of ∼42° C, as determined by DLS.

The observed transformation was found to be fully reversible, albeit with significantly distinct kinetics. Following conversion to native-like oligomer assemblies, the same sample could be readily transformed back to the fibril state through incubation at room temperature (**Fig. 3g**). This process of fibril reassembly occurred over a span of several days, typically reaching completion after 3-4 weeks. Notably, fibrils formed during incubation were markedly longer than those initially isolated from bacterial expression (reaching >1μm in size), with otherwise identical apparent morphology. This reversible transformation process served as the basis for preparing samples for high-resolution cryo-EM analysis, described above.

Remarkably, fibril disassembly could be further repeated, with samples readily converting back to native-like oligomeric structures at an incubation temperature of 42° C (**Fig. 3g**). An intrinsic implication of these observations is that subunit-exchange dynamics, a hallmark feature of native sHSPs, is preserved in the αB-AXA fibril state.

### Disrupting the NT-IXI motif reduces chaperone potency and potentiates client-induced co-aggregation

Previous studies have suggested a role for the NT-IXI motif in αB-crystallin in client binding interactions^38^. To address the potential functional effects of disrupting the NT-IXI motif, we leveraged the temperature dependence of the fibril and native-like oligomer states to assess the impact on chaperone function (**Fig. 4**). Chaperone activity was gauged by monitoring the suppression of light scattering aggregation during the chemically-induced unfolding of a model client, lysozyme^16^. In each experiment, lysozyme (10 μM concentration) was prepared in our standard pH 7.4 buffer, complemented with 150 mM NaCl and 1 mM EDTA, and unfolding of lysozyme was triggered by addition of reducing agent (1 mM TCEP) and monitored by light scattering at 360 nm. This was performed either without chaperone (negative control) or with varying amounts of αB-AXA. For comparison, αB-wt was tested under identical conditions as a positive control.

**Figure 4.**
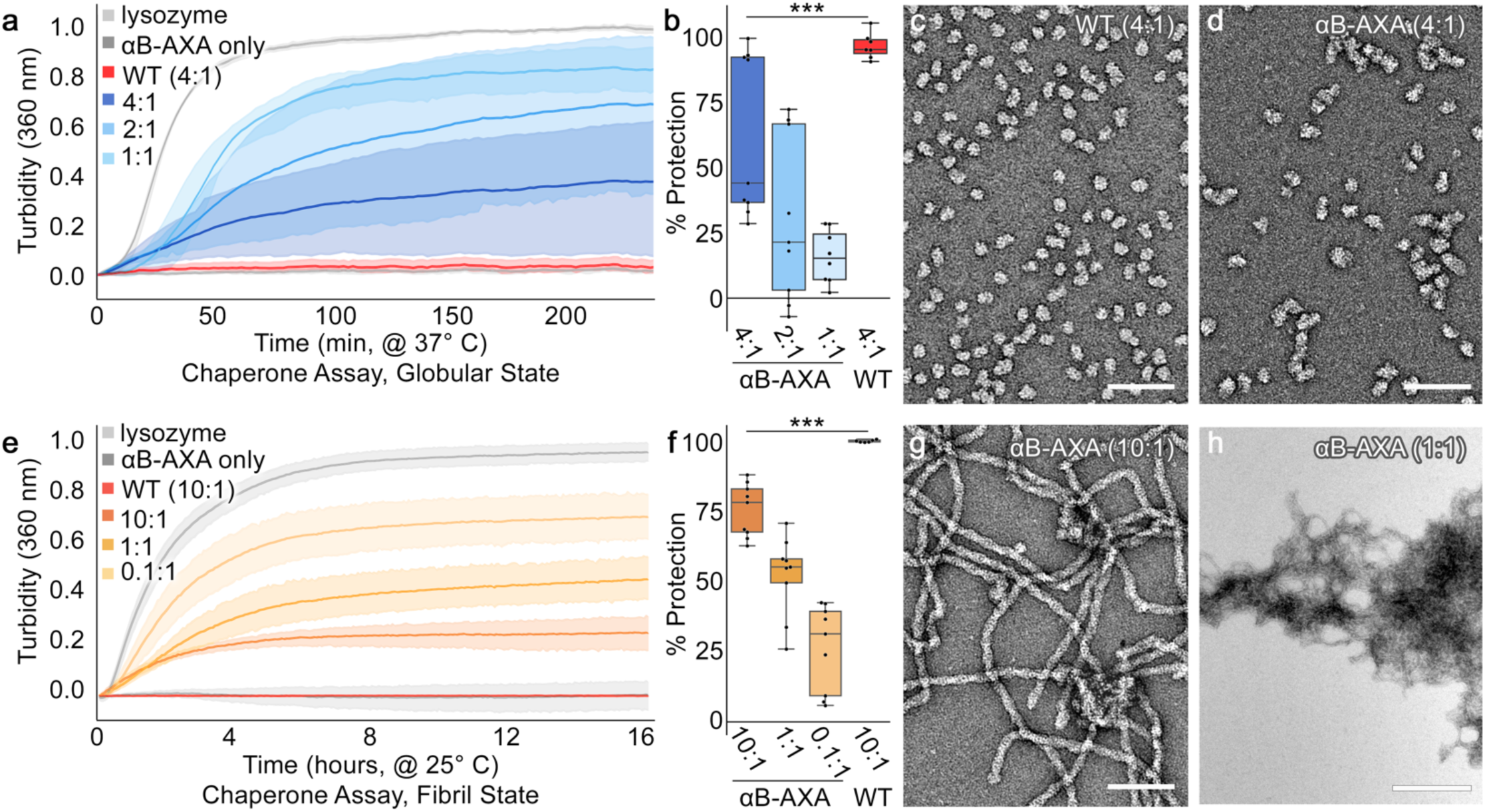
Disruption of the NT-IXI motif results in reduced chaperone activity in both native-like globular and fibril states. **a.** Chaperone assays against unfolding lysozyme client conducted with varying molar ratios of αB-AXA in the native-like globular state (blue traces), as monitored by light scattering at 360 nm at 37° C. Lysozyme-only (10 μM, light grey trace) and αB-wt (40 μM, dark gray trace) were run as positive and negative controls, respectively. αB-AXA prepared at 1:1, 2:1 and 4:1 (chaperone:client) molar ratios. Data are normalized to lysozyme-only conditions. Number of replicates for each experiment (n = 7–9). Standard error of the mean (s.e.m.) shown with semi-transparent shading. **b.** Percent protection summarized in box plot representation and colored as in panel a. Statistical significance (p<0.005; ***). **c, d.** Representative electron micrographs of end-state reactions for the 4:1(chaperone:client) ratios obtained for αB-AXA and αB-wt, respectively. Scale bar = 250 nm. αB-AXA chaperone/client complexes appear as more irregular and elongated particles, as compared to αB-wt under these same conditions. **e.** Chaperone assay of αB-AXA in the fibril state conducted at 25° C (orange-yellow traces). Lysozyme-only (10 μM, light grey trace) and αB-wt (40 μM, dark gray trace) were run as positive and negative controls, respectively. αB-AXA prepared at 0.1:1, 1:1 and 10:1 (chaperone:client) molar ratios. Number of replicates for each experiment (n = 7–9). Standard error of the mean (s.e.m.) shown with semi-transparent shading. **f.** Percent protection summarized in box plot representation and colored as in panel e. Statistical significance (p<0.005; ***). **g, h.** Representative electron micrographs of end-state reactions for the 10:1 and 1:1 (chaperone:client) ratios obtained for αB-AXA fibril state, respectively. Scale bar = 250 and 500 nm in panels g and h, respectively. Increasing client ratios correlate with an increase tangling and/or co-aggregation of αB-AXA fibrils.

To evaluate the NT-AXA variant within the context of the native-like oligomer, samples were fully converted to this state by pre-incubating at 42° C, then sustained in this state at an incubation temperature of 37° C. At this temperature, reduced lysozyme shows robust and consistent aggregation kinetics (**Fig. 4a**, *light grey*). When combined at a 4:1 stoichiometric ratio (40 μM chaperone), αB-wt effectively inhibits light-scattering aggregation (96.7% ± 1.9 protection) (**Fig. 4a**, *red*). At the same 4:1 ratio, the αB-AXA variant in the native-like oligomer state was significantly less effective (62.0% ± 10.3 protection, p<0.005) (**Fig. 4a,b**, *dark blue*). The suppression of aggregation further diminishes at lower chaperone:client ratios, and at a 1:1 ratio the αB-AXA variant displayed only modest protection (16.1% ± 3.6) (**Fig. 4a,b**, *light blue*).

A morphological comparison by EM conducted on the 4:1 reactions highlight the differences in chaperone efficacy between αB-wt and αB-AXA oligomers (**Fig. 4c,d**). We have demonstrated in previous work that under conditions where αB-wt offers complete protection from aggregation, the resultant chaperone-client complexes bear a close resemblance to the apo-state^16^. This finding is corroborated in our current study (**Fig. 4c**). In contrast, αB-AXA/client complexes, when examined at an identical 4:1 stoichiometric ratio, manifest a more irregular and notably elongated morphology (**Fig. 4d**). Such elongated chaperone-client forms have been demonstrated for αB-wt in scenarios where its chaperone capacity is challenged^16^. These observations suggest that while the NT-IXI motif isn’t essential for chaperone activity, its alteration significantly impacts chaperone efficacy, consistent with the proposed role of the NT-IXI in client recognition and/or binding.

Next, to assess chaperone activity of the αB-AXA variant in its helical fibril form, we repeated the aggregation suppression assay at 25° C, a temperature where the fibril structure remains stable. While lysozyme unfolding kinetics decrease considerably at this temperature, they remained robust and consistent, as observed by monitoring light-scattering (**Fig. 4e,** *light grey*). Remarkably, even in its fibril form, αB-AXA retains appreciable chaperone activity, though apparently significantly reduced compared to wildtype. At the highest tested chaperone concentration, 10:1 stoichiometric ratio (chaperone:client), αB-AXA fibrils appear to suppress only 75.4% ± 3.1 of aggregating lysozyme activity (**Fig. 4e,f**, *yellow*). Conversely, αB-wt exhibits nearly total protection under identical conditions (99.9% ± 0.2 protection) (**Fig. 4e,f**, *red*). It’s worth highlighting that at a 1:1 ratio αB-AXA fibrils still offer significant protection, reducing aggregation by 52.1% ± 4.7 (**Fig. 4e,f**, *orange*) and remarkably, at a 0.1:1 ratio a protection of 26.4% ± 5.1 is still present (**Fig. 4e,f**, *dark orange*).

Morphological evaluation of chaperone reactions with αB-AXA fibers unveiled complex behaviors. In scenarios where αB-AXA fibers were most effective (10:1 ratio), fibril morphology largely resembles the apo-state, albeit with the appearance of some induced fibril clustering (**Fig. 4g**). More intriguingly, under higher client ratios (*e.g.,* 1:1 ratio) extensive fibril clustering is observed (**Fig. 4h**). EM images showed massive fibril tangles, some spanning up to 2-3 μM in diameter. Such large aggregates would be sufficient to contribute to light scattering at 360 nm, used for monitoring chaperone activity, complicating interpretation of the aggregation suppression assay. Nevertheless, it is evident that client-interaction with the αB-AXA fibrils induces co-aggregation in this system. A possible interpretation of these results is that the unfolded client gets recognized and simultaneously tethered to multiple fibrils, triggering fibril cross-linking and co-aggregation. However, as the unfolded client is not resolved in the EM images, further studies are needed to confirm this proposed model.

## DISCUSSION

Our findings underscore the pivotal structural and functional role of the NT-IXI motif within the αB-crystallin sHSP chaperone. By eliminating the expected competitive binding between the NT-IXI and CT-IXI at the ACD β4/8 hydrophobic groove, we observed a marked shift in the equilibrium-state assembly of αB-crystallin towards a novel fibril state. This fibril formation, albeit unexpected, finds parallels in a distant sHSP counterpart found in Salmonella bacterium: AgsA. In AgsA, this fibril state is heat-inducible and believed to guard against irreversible aggregation, and reverses upon introduction of destabilized client proteins^54^. Such a mechanism in sHSPs hasn’t been proposed for eukaryotic organisms. Rather, it is expected that the fibril state of αB-crystallin observed in our study stems from the perturbation to the NT-IXI motif.

Despite the profound impact on quaternary assembly induced by the NT-AXA variant, many key features expected for sHSPs are preserved that help to both validate and significantly deepen our understanding for how sHSPs assemble into dynamic oligomers. The ACD dimer forms the fundamental building block in the assembly in both the fibril state and the native oligomeric states^41^. Previous structural investigations on truncated αB-crystallin constructs, which contained only the ACD domain, identified multiple alignment registers at the dimerization interface (AP_I_, AP_II_ and AP_III_). Solid-state NMR analyses of the full-length αB-crystallin oligomer primarily detected the AP_II_ register^39^, similar to our analysis of the fibril state. While there’s potential for the presence of minor populations with alternative registers in the αB-AXA fibril state, our rigorous 3D classification attempts did not provide any evidence for this.

The β5/6 loop of the ACD has exhibited multiple conformational states, which are broadly categorized as the up or down conformation^31^. In the αB-AXA fibril state, the upward conformation is resolved, and stabilized by salt-bridge interactions involving highly conserved D109, R116 and R120, with D109 forming a key interaction with R120 across the dimer interface (see **Supplemental Fig. 5**). Genetic mutations affecting D109 and R120 have been linked to familial cataract and various myopathies^55–58^. These mutations have been shown to alter the structural state of the αB-crystallin oligomer and have a negative impact on its chaperone function^59–62^, underscoring a critical importance to maintaining the structural-functional integrity of the holo-complex.

Another site of variance across ACD dimers is the presence of a short anti-parallel β2 strand. This feature is not always resolved in various X-ray and NMR structures of the αB-crystallin ACD^27,36,39^, suggesting an intrinsic level of conformational flexibility. Remarkably, this feature is not symmetrically resolved across the ACD dimer in the fibril assembly, and only observed for subunits along the external seam. In this context, β2 overlays the bound NT region, presumably contributing to the localized stability. Consistent with the notion of β2 serving as a dynamic component of αB-crystallin assembly, this region of the cryo-EM map was considerably less-well resolved as compared to other regions of the ACD.

The CTD of αB-crystallin is also understood to be dynamic, presenting a challenge to assessing the details of its interaction with the ACD in the context of an oligomeric assembly. With the NT-IXI motif ablated, the CTD forms a persistent interaction with the ACD β4/8 hydrophobic groove that could be well-resolved in the fibril state. Intrinsic flexibility within the CTD linker appears to provide two critical functions. First, it allows for multiple conformational states, enabling distinct domain-swap interactions between different neighboring ACD dimers. This stable interaction is accommodated by the hydrophobic knob-in-hole interactions between the conserved CT-IXI motif and the ACD edge-groove, and the palindromic sequence around the CT-IXI motif permits a bi-directional orientation of the CTD within the ACD groove. Secondly, the flexibility of the CTD linker connecting the ACD to the IXI motif appears to impart long-range structural plasticity to the high-order assembly. Similar flexibility within the CTD has been suggested in other sHSP assemblies^63,64^, and is likely critical to the capacity for adopting various oligomeric states and for accommodating different clients.

The ability of the αB-AXA variant to rapidly transform between the fibril and native-like oligomer states is evidence of underlying subunit exchange dynamics, considered to be a hallmark of sHSP structure, polydispersity, and chaperone function^65^. It was postulated that an interplay of competitive binding between the NT-IXI and CT-IXI for ACD β4/8 groove binding influences the polydispersity of αB-crystallin^36^. In the context of the αB-AXA variant, the CT-IXI has uninhibited access the ACD binding site, favoring a more monodispersed structure, which is the elongated fibril state. At elevated temperatures, thermal dynamics could amplify the CT-IXI’s dissociation rate, simulating the rapid-exchange dynamics experienced under native conditions of competition with the NT-IXI, and promoting a shift toward the native-like globular state (**Fig. 5**). While this explanation seems parsimonious to these data, future studies will be needed to further investigate and confirm this proposed model.

**Figure 5.**
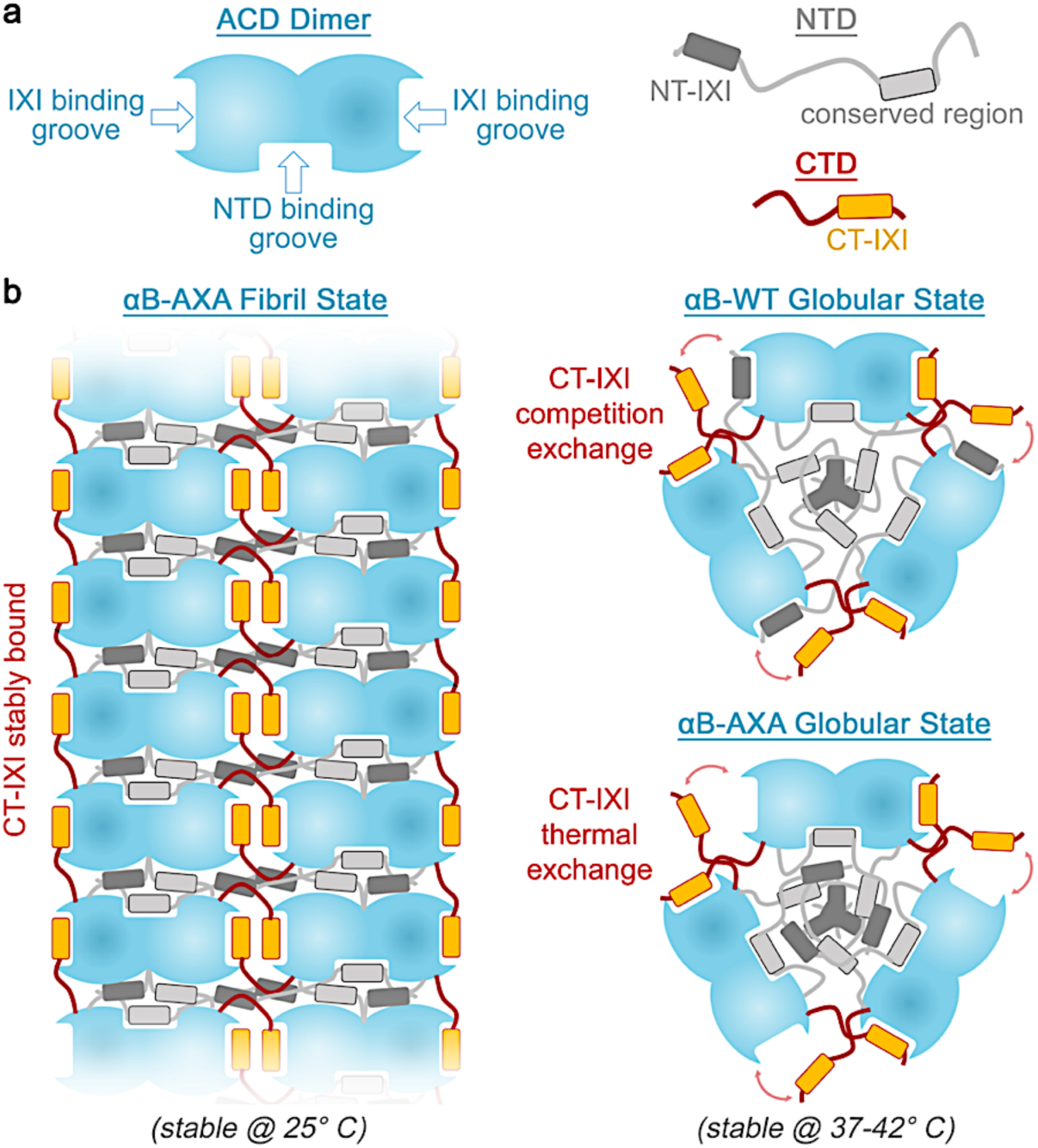
Overview and proposed structural organization of αB-crystallin in fibrillar and globular states. **a.** Illustration depicting the αB-crystallin ACD dimer building block (blue) that establishes two hydrophobic IXI-motif binding grooves and a putative NTD binding groove at the dimer interface. The NTD (grey) comprises two conserved regions, the NT-IXI motif (dark grey box) and a conserved hydrophobic region (light grey box). The CTD (red) contains the conserved CT-IXI motif (orange box). **b.** When the NT-IXI is ablated (αB-AXA), the CT-IXI can form persistent and stable interactions with the IXI binding grooves in the ACD, promoting elongated fibril assembly. In this state, the NTD is buried within the interior cavity of the assembly. The NTD binding groove of the ACD is occupied and is presumed to correspond to the conserved region of the NTD. Upon heating the αB-AXA construct from 25°C to 37-42°C, the structure transitions into a globular state resembling wild-type αB-crystallin (αB-WT). This transformation is proposed to result from thermal energy causing rapid exchange of the CT-IXI with the ACD IXI-binding groove, mimicking the exchange mechanism proposed in αB-WT, where both the CT-IXI and NT-IXI undergo competition exchange for a limited number of binding grooves in the ACD.

In the αB-AXA fibril morphology, the NTD occupies the internal cavity of the assembly, adopting a quasi-ordered (or partially disordered) state. This aligns with earlier cryo-EM studies of full-length αB-crystallin, where the NTD was either weakly defined within the interior of the cage-like assembly or not discernible at all^13,40,41,48^. Similarly, prior solid-state NMR studies on αB-crystallin indicated the dynamic nature of the NTD, suggesting an ability to adopt multiple conformational states^36,39^. A segment of the NTD was partially resolved in the αB-AXA fibril structure and stabilized by binding along an interior groove formed by the ACD dimer interface (**Fig. 5**). This interaction appears further reinforced by extensive contact with β2, that forms a ‘lid’ that effectively buries the NTD within the ACD dimer interface. Although the local resolution wasn’t sufficient to confidently assign this NTD peptide region, its characteristics appear consistent with the conserved hydrophobic region implicated in similar interaction with the ACD in related sHSPs^36,51,66^ (**Fig. 2d**). A recent crystal structure of the HSPB2/3 heteromeric complex also resolved a density lining the interior ACD groove, which was also attributed to the NTD but could not be confidently assigned due to poor resolution^67^.

The interaction between the NTD and ACD might have additional functional implications beyond its contribution to architectural stability and high-order assembly^27,68^. Particularly noteworthy are the sections of the ACD involved in binding the NTD peptide that have been implicated in client recognition and chaperone activity of αB-crystallin^69^, including the mini-chaperone peptide region (residues 73–92)^70^. Under basal-state conditions, it’s plausible that the NTD-ACD interaction serves to conceal this client-recognition site. Genetic removal of this region of the NTD in αB-crystallin not only results in significant loss of high-order assembly and structural stability, but also increases surface hydrophobicity and enhances chaperone activity^71^. Several phosphorylation sites are also situated within the NTD that are believed to trigger chaperone activation and disrupt the oligomeric assembly^21^. Given this, it’s reasonable to speculate that phosphorylation of the NTD may effectively disrupt this ACD interaction, thereby causing the oligomer to destabilize and unmasking of the client-binding site, ultimately resulting in chaperone activation.

The diminished chaperone capacity of the αB-AXA variant within its native-like globular state aligns with the anticipated role of the NT-IXI motif in client recognition and/or binding, as a previous NMR study identified this motif as contributing to interactions formed with the bound client, lysozyme^38^. It is also possible that NT-AXA variant exhibits altered subunit exchange kinetics that could contribute to the augmented chaperone activity^72^. Notably, the αB-AXA variant still exhibits chaperone function in its fibril state, yet it also potentiated the formation of expansive chaperone-client co-aggregates. This behavior may rationalize the high-degree of conservation of the NT-IXI motif in αB-crystallin, as it suggests such fibrils forms may be detrimental to physiological role in the cell, for example in the context of the eye lens, where such co-aggregates could potentially lead to cataract formation.

The physiological implications of the αB-AXA fibril structure remain uncertain; however, we posit that it may highlight αB-crystallin’s tendency to form highly elongated structures integral to its chaperone function. We recently described a client-driven elongation mechanism shared by both αA- and αB-crystallin^16^ (see also **Fig. 4d**). Given that the NT-IXI motif is believed to play a role in client binding, such engagement could effectively sequester the NT-IXI and eliminate its competition with the CT-IXI for ACD binding. Consequently, the mechanistic underpinnings enabling the highly elongated morphologies of αB-crystallin induced upon client binding may be reflected in the αB-AXA phenotype. Future studies aimed at further disentangling the complex dynamics and chaperone mechanism of the sHSPs will be needed to fully address this hypothesis and other new questions that have been opened by this work.

## ACKNOWLEDGEMENTS

We thank Adam Miller and Dr. Kirsten Lampi for helpful discussions. We are grateful to the staff at the OHSU Multiscale Microscopy Core and Advanced Computing center, and to the Pacific Northwest Center for CryoEM (supported by NIH Grant U24GM129547) and accessed through EMSL (grid.436923.9). The research was funded by NIH grants R01EY030987 and R35GM124779 (to S.L.R.) and fellowship F31EY033230 (to R.M.).

## AUTHOR CONTRIBUTIONS

R.M. performed the experiments and analyzed the data; S.L.R. and R.M. conceived of the study and prepared the manuscript.

## METHODS

### NT-IXI motif identification, construct design and protein purification

The NT-IXI motif was identified by aligning the amino acid sequences of α-crystallins and other related small heat shock proteins. To generate the NT-AXA variant, site-directed mutagenesis was performed on a pET plasmid containing the human αB-crystallin gene, where residues I3 and I5 were replaced with alanine using the Quick-Change Lightning kit following the manufacturer’s instructions (Agilent). Following mutagenesis, modified plasmids were introduced into XL-10 Gold cells for amplification. Plasmid DNA was then extracted using Promega miniprep kits and sequenced to confirm the introduction of the desired mutations.

For protein expression, DNA plasmids were transformed into BL21 *E. coli* cells, by heat shock at 42° C for 45 seconds. Cells were cultured in LB media at 37° C under ampicillin control until they reached an optical density (OD) of 0.5. The culture temperature was then reduced to 18° C, followed by induction with 1 mM IPTG for protein expression overnight (∼18 hours). Cells were harvested by centrifugation and the pellet was re-suspended in 4 mL of lysis buffer containing 20 mM Tris pH 8, 1 mM EDTA (4 mL per gram of pellet) and stored at −80° C for subsequent protein purification.

Cell suspensions were thawed and treated with 0.1 mM DTT and PMSF, and lysis was carried out by sonication (Fisher Sonic Dismembrator), with settings at 70% intensity and pulse times of 30 seconds, for a total duration of 6 minutes. After sonication, an additional 0.1 mM PMSF was added to the lysate to prevent proteolytic degradation. The lysate was then ultra-centrifuged at 147,000 x g for 30 minutes at 4° C to remove insoluble cellular components. The supernatant was retained and further treated with 20 units of DNase I to degrade residual DNA. Following DNase treatment, the lysate was filtered through a 0.45 μm filter.

For protein purification, the clarified lysate was applied to a Sephacryl 300 column (Cytiva Life Sciences) that had been pre-equilibrated with a salt-free size-exclusion chromatography (SEC) buffer (20 mM Tris at pH 8.0, 1 mM EDTA, and 0.1 mM DTT). Fractions containing the αB-crystallin were identified by SDS-PAGE, pooled, and supplemented with 0.1 mM DTT. Pooled fractions were then loaded onto a MonoQ ion-exchange column (Sigma-Aldrich) and eluted across a salt gradient of 0.5 M NaCl. Elution peaks containing αB-crystallin were collected, pooled, and subsequently dialyzed with a 3.5 kDa molecular weight cut-off (m.w.c.o.) membrane (SnakeSkin, Thermo Scientific) against the final reaction buffer (20 mM HEPES, 100 mM NaCl, 1 mM EDTA, pH 7.4).

### Temperature dependence assays by DLS and EM

To elucidate the transitions between fibrillar and globular assemblies, samples were incubated at varying temperature and monitored by dynamic light scattering (DLS) and electron microscopy (EM). DLS assays were performed on a Wyatt DynaPro III instrument, with the resulting data processed using the DYNAMIC 6 software package. For initial characterization, protein samples were subjected to a range of temperatures, including 4° C, 25° C, and 42° C. For temperature ramp assays, samples were started at 25° C and progressively heated to 60° C (0.33°/minute), beyond which point irreversible aggregation was observed. Morphological transitions were detected by DLS and subsequently confirmed by EM.

Extended incubation at 25° C produced samples with long, well-ordered fibrils. Full conversion to the fibril state was observed after incubations of up to 4 weeks, as confirmed by EM. Following this incubation period, samples used for structural analysis by cryo-EM underwent an additional step of dialysis to remove trace amounts of unconverted globular oligomers using a dialysis membrane with a 1 MDa m.w.c.o. (Repligen Spectra/Por, Spectrum Chemicals) and stored at 4° C.

### Negative Stain Electron Microscopy

Negatively stained samples were prepared similarly for all specimens for EM analysis by placing 3 µL drops of the protein solution onto carbon-coated copper mesh grids at approximate monomer concentrations of 720 nM. After a brief incubation period, excess solution was blotted away on filter paper, washed three times with water, stained with 0.75% uranyl formate (SPI-Chem), and subsequently dried under a laminar flow. Electron micrographs were obtained using a 120 KeV Tecnai TEM (FEI) with a BMEagel detector recorded at a nominal 49,000 x magnification with calibrated pixel sizes of 4.37 Å pixel^-1^.

Initial structural analysis of αB-AXA fibrils was performed on EM images of negatively stained specimens. Preliminary fibrillar crossover dimensions were obtained by direct measurements using the Fiji software^73^. For initial 2D and 3D analysis, individual filament segments were selected using the ‘helixboxer’ tool in EMAN2^74^ and extracted using a box size of 88 pixels and a 90% box overlap. Extracted particles were imported into Relion^75^ for class averaging and a *de novo* initial model was generated in EMAN2.

For cage-like assemblies of αB-AXA, particles were automatically picked using the threshold picking tool in EMAN2 and extracted using a box size of 84 pixels before being imported into Relion for 2D class averaging.

### Cryo-Electron Microscopy

αB-AXA samples that had undergone elongation at room temperature for about four weeks were prepared for cryo-EM analysis. Elongated samples were dialyzed with a 1 MDa m.w.c.o. membrane, to remove any residual globular assemblies. For cryo-grid preparation, 3 µL drops of the sample were applied to R2/1 Quantifoil grids at a concentration of 0.5 mg/mL and plunge frozen using a Vitrobot Mk III autoplunger (FEI). Image acquisition was conducted on a Titan Krios G3 (Thermo-Fisher), equipped with a K3 direct electron detector (Gatan). The completed dataset was obtained using automated data collection routines with SerialEM^76^, resulting in 10,654 micrographs with a defocus range of 0.5 to 2 µm. Movies were recorded with a physical pixel size of 0.788 Å/pixel (0.394 Å/pixel super-resolution) and a total electron dose of 50 e^-^/pixel.

For image processing, electron micrographs were imported into cyoSPARC^46^ in two subsets (containing 4585 and 6069 micrographs), where they were motion corrected and CTF fit. The Filament Tracer module in cryoSPARC was used to pick particles from a random subset of 10 micrographs with fibril diameter 160 Å and a segment overlap of 0.2 x diameter, which were then averaged using 2D classification to generate initial templates for subsequent particle picking. This was then applied to an extended subset of 100 micrographs, leading to a total of 27,990 segments. These segments were then classified to generate a refined template and applied to the entire dataset, yielding 2.2 million segments with a box size of 400 pixels.

Several rounds of 2D classification were applied to the full dataset with a box size of 400 pixels to first remove non-protein and low-quality segments. Later classification steps sought to remove segments with poor alignment near the edge of the box. A total of 1,406,353 well-aligned segments were extracted using a box size of 288 pixels and used to generate an initial model from *ab initio refinement* with a total of 4 classes. The highest quality class had initial helical parameters of 31° twist and 39 Å rise. Helical parameters were iteratively improved through multiple rounds of helical refinement with D2 symmetry applied, which resulted in final helical parameters of 31.6° twist and a rise of 39.9 Å at a resolution of ∼4.1 Å. This particle stack was then masked to isolate the internal D2 symmetric unit and subjected to local refinement with unchanged box size, reaching a final resolution of 3.4 Å (gold-standard FSC; see **Extended Data Table 1** and **Extended Data Figure 3**). Additional workflows were extensively assayed incorporating 3D classification and local refinement in cryoSPARC (as well as in Relion^77^), but these did not lead to appreciable improvement of resolved features.

### Model Building

Atomic models for the asymmetric unit were built starting with the previously reported crystal structure of the human αB-crystallin ACD dimer (PDB 2WJ7)^27^, which was fitted into the unsharpened cryo-EM map obtained from local refinement. The model of the ACD dimer was first rigidly fit into the map with ChimeraX^78^ and then flexibly fit using ISOLDE^79^. The CT-loop at the inner seam and CT-IXI motifs at both the inner- and outer-seam were sufficiently resolved to be manually built in COOT directly from the local refinement map. The CT-loop along the outer seam though was lost in the local refinement job. Model building in this region required the alignment of several ACD models in a lower resolution map containing multiple rungs and the connecting loops were built in COOT and fit flexibly with ISOLDE. This preliminary model was then refined through iterations of refinement in Phenix^80^ and adjustments in both ISOLDE and COOT^81^ until refinement statistics converged, as judged by Molprobity^82^ (see **Extended Data Table 1**).

### 2D and 3D Variability Analysis

Conformational variability of fibrils observed in 2D classification results obtained from cryoSPARC was assessed by hand in Fiji^73^. A total of eight representative classes were chosen based on presence of bend angle (n = 4) or crossover distance (n = 4), as examples of the variability observed from the 2D classification results. Crossover distances in 2D classes was determined by measuring distance between successive cross-over points observed in projection, defined as either the neighboring narrowest or broadest regions of the fibril. Bend angles were measured using the angle measurement tool in Fiji, covering the three crossovers that were visible in a single 2D class.

To further assess the intrinsic variability of the helical assembly, 3D variability analysis (3DVA) was conducted using cryoSPARC^53^. Initially, particles were extracted with a box size of 800 pixels to capture the variance along the length of the fibril. This analysis revealed distinct principal components describing both stretching and bending modes of variance. To enhance the quality of the volume series for the bending mode, the “intermediates output type” was used, which reconstructs the volume for every individual frame rather than interpolating from a central point. This method yielded a more well-defined volume series (**Supplemental Movie 2**).

The fibril’s stretching mode was less obvious in this initial analysis. To enhance this feature, particles were culled from the 2D classification process to remove classes that exhibited bent conformations and performed 3DVA on the resultant particle stack. This approach revealed distinct principal components describing the stretching mode with increased magnitude over the previous approach. However, in this case, the density of the volume series remained poor near the edges of the box. To address this issue, particles from either end of the volume series were selected and used to reconstruct the endpoints of the component. The resulting linear interpolation between the two volumes resulted in a higher quality volume series (**Supplemental Movie 2**).

### Chaperone assays

Chaperone assays were carried out by monitoring the suppression of light scattering caused by chemically induced aggregation of the model client, lysozyme (Sigma, mass-spec grade). For all assays, lysozyme was prepared at a final concentration of 10 μM in a reaction buffer of 20 mM HEPES, 100 mM NaCl and 1 mM EDTA (pH 7.4). TCEP was added to a final concentration of 1 mM to induce the aggregation of lysozyme, which was tracked by an increase of turbidity at 360 nm (Tecan Infinite 200 Pro). All assays included reactions containing lysozyme only (negative control) and αB-wt (positive control).

For assays conducted on αB-AXA in the native-like morphology, samples were fully converted to the globular state by incubating at 42° C for ∼18 hours. Full conversion of these samples was confirmed by DLS and EM. Chaperone assays were conducted at 37° C and light scattering was monitored for 4 hours, at which point the reactions had reached steady-state. αB-AXA assemblies were assayed against 10 µM lysozyme over a range of concentrations (10 – 40 µM). Assays conducted using αB-wt were performed under these same conditions. All assays were performed in replicate. Chaperone activity was calculated from the turbidity traces and converted to percent protection versus lysozyme-only conditions.

For assays conducted on αB-AXA fibrils, protein stocks of 3 mg/mL were elongated at 25° C and dialyzed overnight in reaction buffer using a 1MDa m.w.c.o. membrane to remove residual native-like assemblies (as confirmed by EM). Chaperone assays for these samples were conducted at 25° C to maintain stability of the fibril assembly. Light scattering was monitored for 18 hours to account for slower aggregation kinetics of lysozyme at these reduced temperatures. αB-AXA fibrils were assayed against 10 µM lysozyme over a range of concentrations (1 – 100 µM). EM grids were prepared on negatively stained samples of the end-point reactions and imaged as described above.

### Statistical Analysis

All chaperone assays were run in replicate (n = 7–9). Raw turbidity data from chaperone assays were processed by first min-max normalizing. Percent protection was determined by the percent reduction in turbidity compared to the lysozyme-only control samples and significance was evaluated by two-sample t-tests.

### Figure Preparation

Structural models and cryo-EM density maps were visualized and prepared for presentation using ChimeraX^78^. Cartoon illustrations were inspired by previous work by Reinle *et al*^3^, and prepared in PowerPoint. Final figures were composed in Photoshop.

### AI-assisted technologies

During the preparation of this work the authors used ChatGPT to help revise portions of the text to improve readability. After using this tool, the authors reviewed and edited the content as needed and take full responsibility for the content of the publication.

### Data Availability

Cryo-EM density maps have been deposited to the Electron Microscopy Data Bank (EMD-XXXX). Coordinates for atomic models have been deposited to the Protein Data Bank (XXXX). The original multi-frame micrographs have been deposited to EMPIAR (EMPIAR-XXXXX).

## EXTENDED DATA (TABLES, FIGURES AND LEGENDS)

**Extended Data Table 1.**
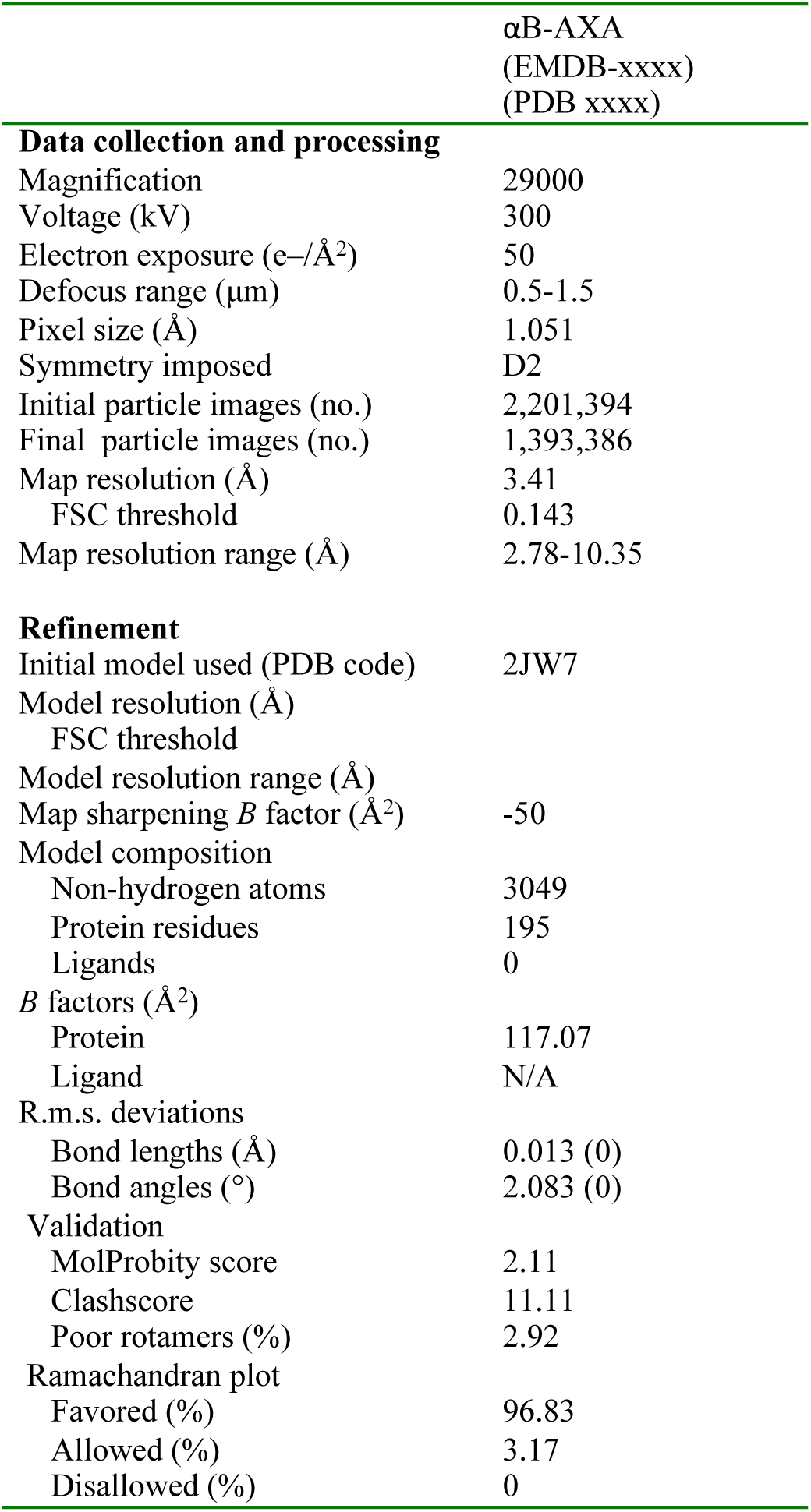
Cryo-EM data collection, refinement, and validation statistics.

| | αB-AXA<br>(EMDB-xxxx)<br>(PDB xxxx) |
| --- | --- |
| **Data collection and processing** |  |
| Magnification | 29000 |
| Voltage (kV) | 300 |
| Electron exposure (e-/Å<sup>2</sup>) | 50 |
| Defocus range (μm) | 0.5-1.5 |
| Pixel size (Å) | 1.051 |
| Symmetry imposed | D2 |
| Initial particle images (no.) | 2,201,394 |
| Final particle images (no.) | 1,393,386 |
| Map resolution (Å) | 3.41 |
| FSC threshold | 0.143 |
| Map resolution range (Å) | 2.78-10.35 |
| **Refinement** |  |
| Initial model used (PDB code) | 2JW7 |
| Model resolution (Å) |  |
| FSC threshold |  |
| Model resolution range (Å) |  |
| Map sharpening *B* factor (Å<sup>2</sup>) | -50 |
| Model composition |  |
| Non-hydrogen atoms | 3049 |
| Protein residues | 195 |
| Ligands | 0 |
| *B* factors (Å<sup>2</sup>) |  |
| Protein | 117.07 |
| Ligand | N/A |
| R.m.s. deviations |  |
| Bond lengths (Å) | 0.013 (0) |
| Bond angles (°) | 2.083 (0) |
| Validation |  |
| MolProbity score | 2.11 |
| Clashscore | 11.11 |
| Poor rotamers (%) | 2.92 |
| Ramachandran plot |  |
| Favored (%) | 96.83 |
| Allowed (%) | 3.17 |
| Disallowed (%) | 0 |

**Extended Data Figure 1.**
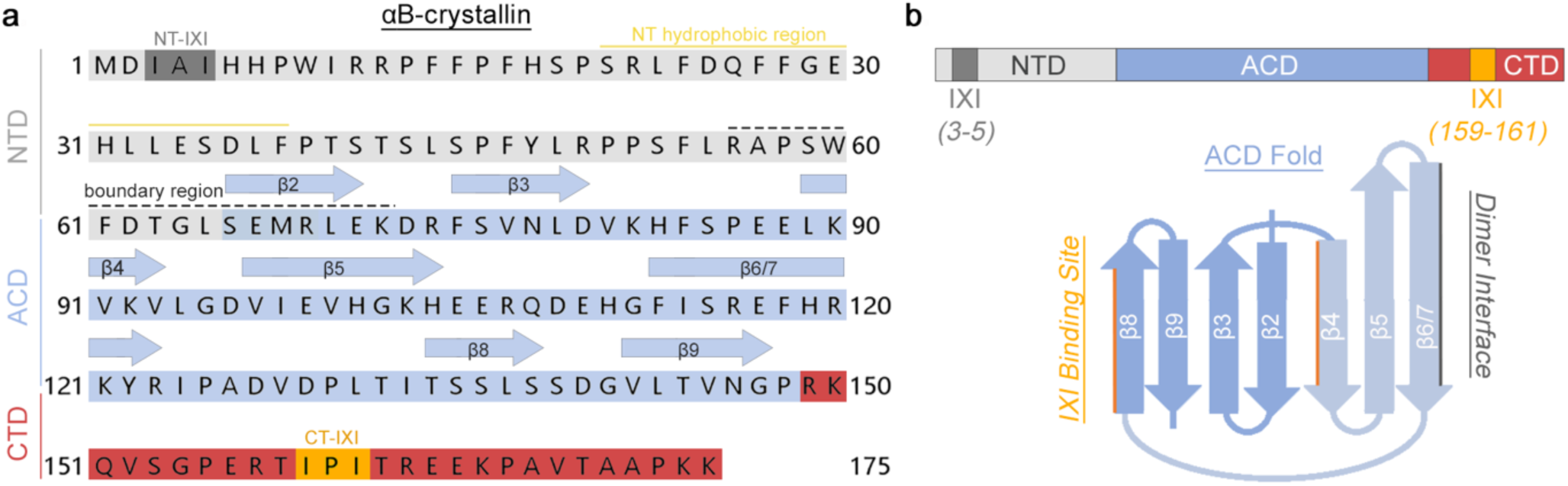
Annotated primary sequence and structural topology of αB-crystallin. **a.** Primary amino acid sequence of human αB-crystallin. Structure features colored and annotated as: N-terminal domain (NTD) (light gray), α-crystallin domain (ACD) (light blue), C-terminal domain (CTD) (red), NT-IXI motif (dark gray), CT-IXI motif (orange), conserved hydrophobic region of the NTD (yellow line), NTD boundary region (black dashed line). ACD secondary structural elements for β-strand 2-9 indicated by blue arrows. **b.** Linear (top) and secondary structural topology (bottom), colored as in panel a. Secondary structural components involved in forming the ACD dimer interface (gray line) and CT-IXI binding site (orange line) are indicated. Blue shading indicates the two β-sheets formed by the ACD fold.

**Extended Data Figure 2.**
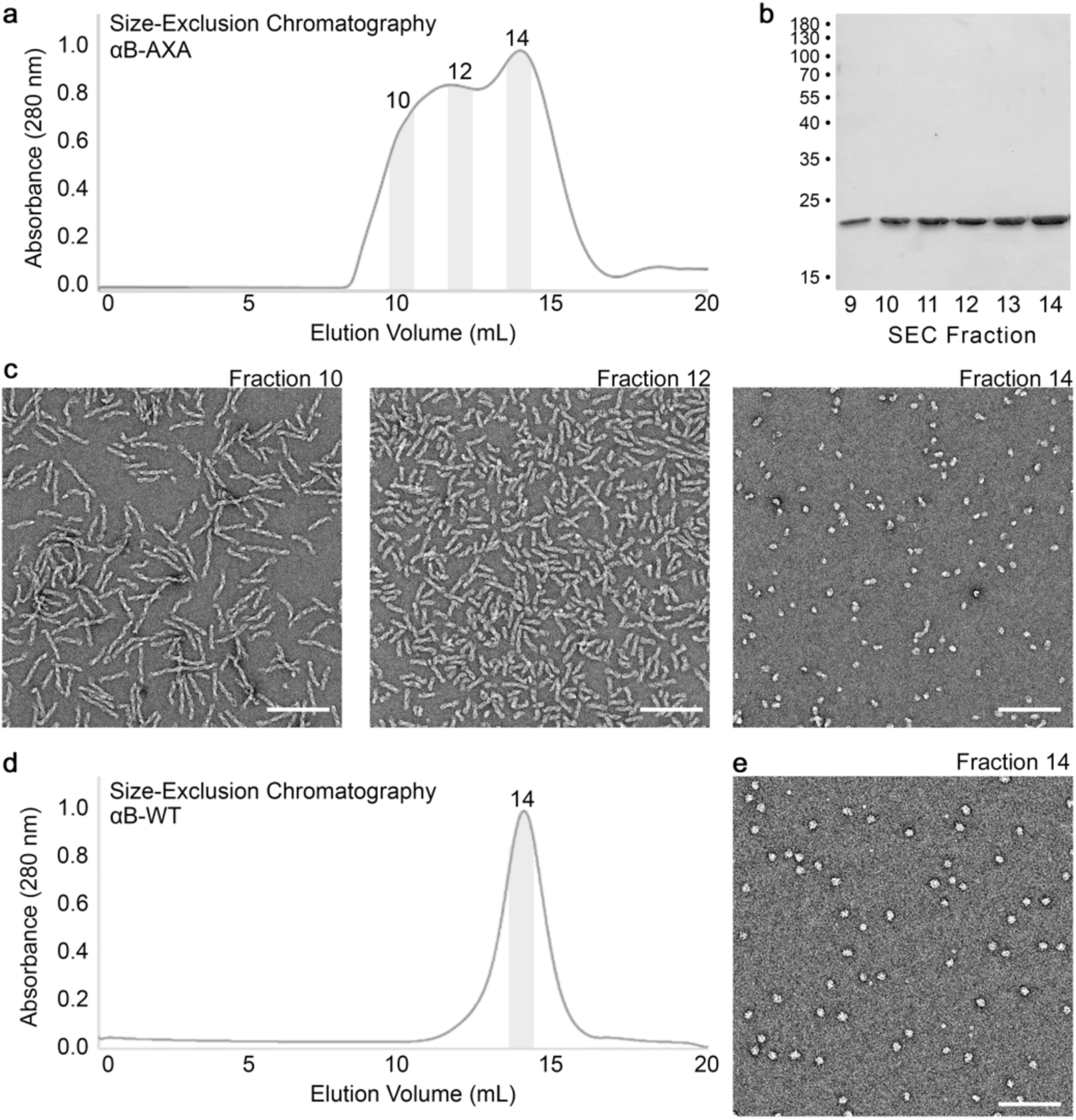
Purification of wildtype αB-crystallin and NT-AXA variant. **a.** Size exclusion chromatography (SEC) elution profile monitored by absorbance at 280 nm for αB-AXA, shows a main peak centered at ∼14 mL with a broad left-hand plateau (superose-6, 24 mL column). **b.** SDS-PAGE analysis of SEC elution fractions (9 – 14) for αB-AXA showing a single band at ∼20 kD present in all fractions. **c.** Electron micrographs of negatively stained αB-AXA of selected SEC elution fractions, labeled and indicated by grey shading in panel a (scale bar 100 nm). **d.** Size exclusion chromatography elution profile of αB-wt with characteristic peak centered at ∼14 mL (superose-6, 24 mL column). **e.** Electron micrograph of negatively stained αB-wt particles (scale bar 100 nm). Selected SEC fraction is labeled and indicated by grey shading in panel d.

**Extended Data Figure 3.**
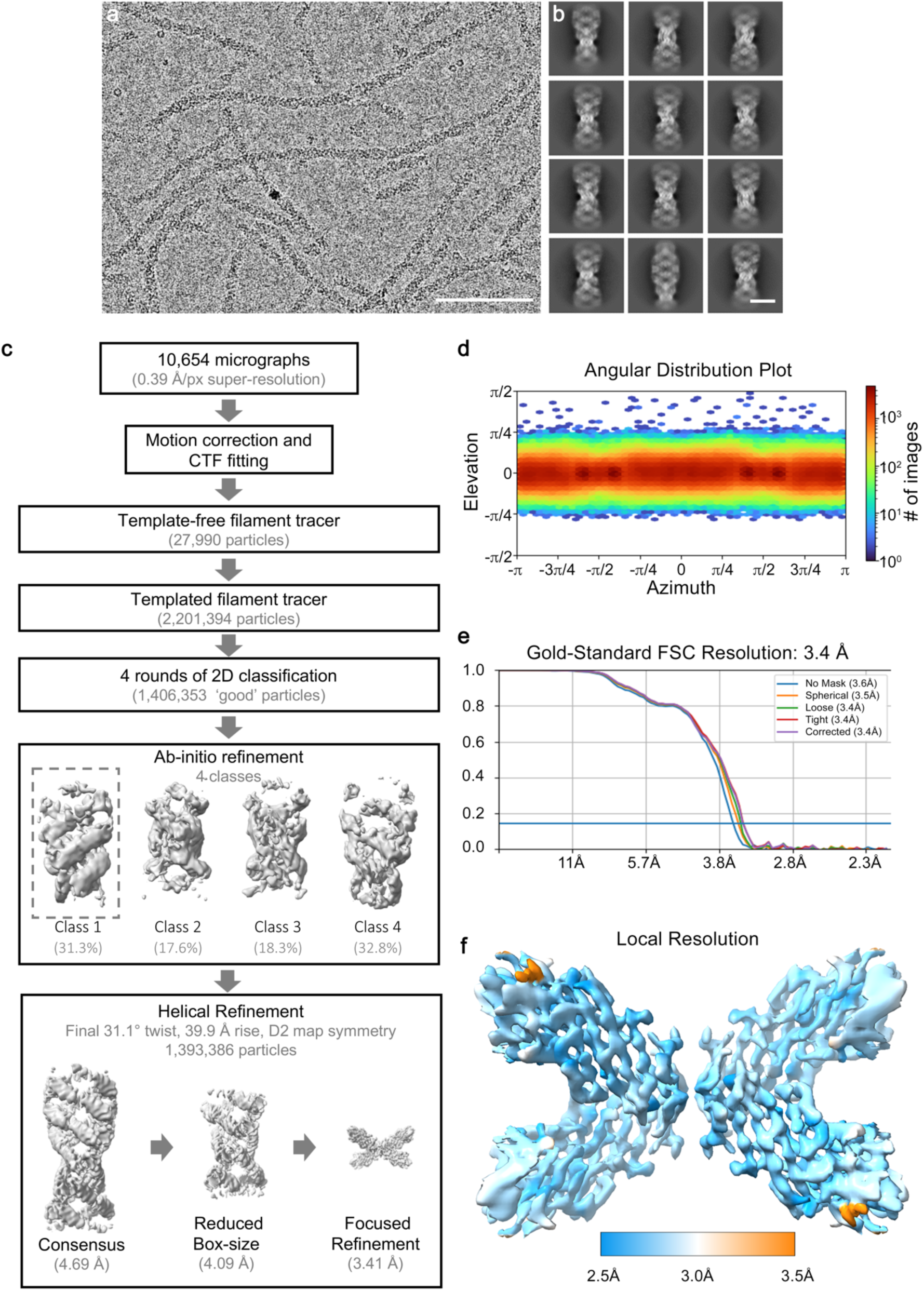
Overview of cryo-EM image processing workflow. **a.** Representative micrograph of αB-AXA fibrils recorded by cryo-EM and **b.** resulting 2D class averages of isolated fibril segments. Scale = 100nm and 10 nm, respectively. **c.** Schematic of the cryo-EM image processing workflow. A total of 10,654 micrographs were collected in an automated fashion using SerialEM on a 300 kV Titan Krios equipped with a K3 direct detector with a physical pixel size of 0.788 Å (binned 0.394 Å super-resolution). Movies were corrected for drift and CTF fit in cryoSPARC^46^. A set of 27,990 segments were picked in a template-free manner using the Filament Tracer. These segments were then subjected to 2D classification to generate templates for templated particle picking, which yielded a set 2,201,394 fibril segments. 2D classification was used to clean this particle stack to 1,406,353 particles utilizing a large box size (400 px) to isolate segments with apparent long-range order. Particles were then re-extracted with a smaller 288 pixel box size for *ab initio* refinement with 4 classes to generate an initial models. A single class containing approximately 31% of the particle showed reasonable helical symmetry and domain features and was selected for subsequent helical refinement using the entire particle stack and applied D2 symmetry, resulting in a resolution of 4.1 Å. **b.** Particles were then masked for local refinement of the central D2-symmetric unit resulting in a final global resolution of the cryo-EM map of 3.4 Å. **d.** Angular distribution plot of refined particles. **e.** Gold-Standard Fourier-Shell Correlation (FSC) analysis generated in cryoSPARC demonstrating a global resolution of 3.4 Å. **f.** Local resolution assessment of the refined cryo-EM map (range of 2.5 to 3.5 Å, blue – white – orange).

**Extended Data Figure 4.**
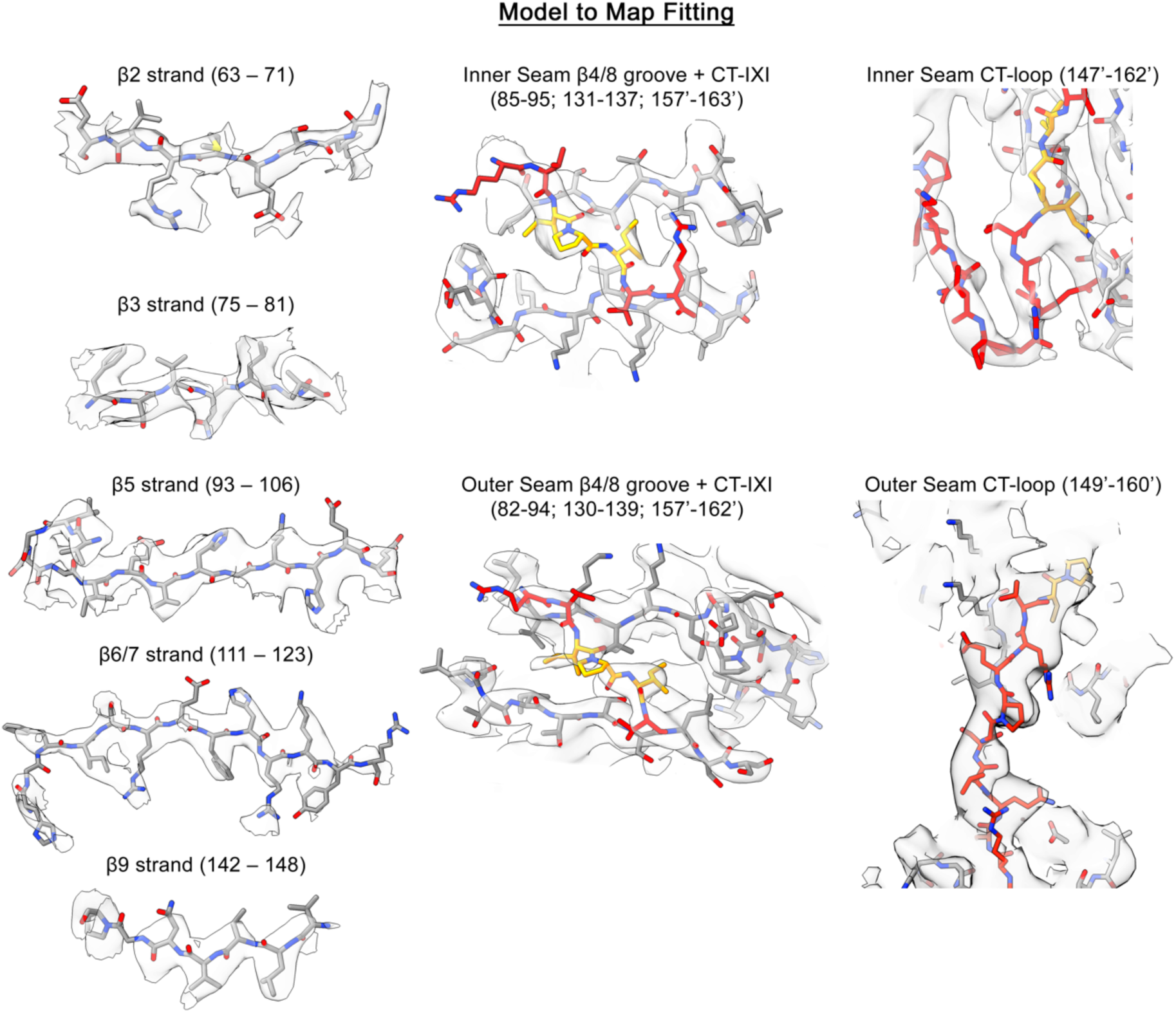
Fit of αB-AXA atomic model to the cryo-EM density. Representative views of the αB-AXA model (stick representation) fit to cryo-EM density. The cryo-EM map was segmented and shown in transparency for clarity. The atomic model is colored by heteroatom (oxygen – red, nitrogen – blue), with base color for carbon atoms within the ACD colored in grey, CTD colored in red and CT-IXI motif colored in orange.

**Extended Data Figure 5.**
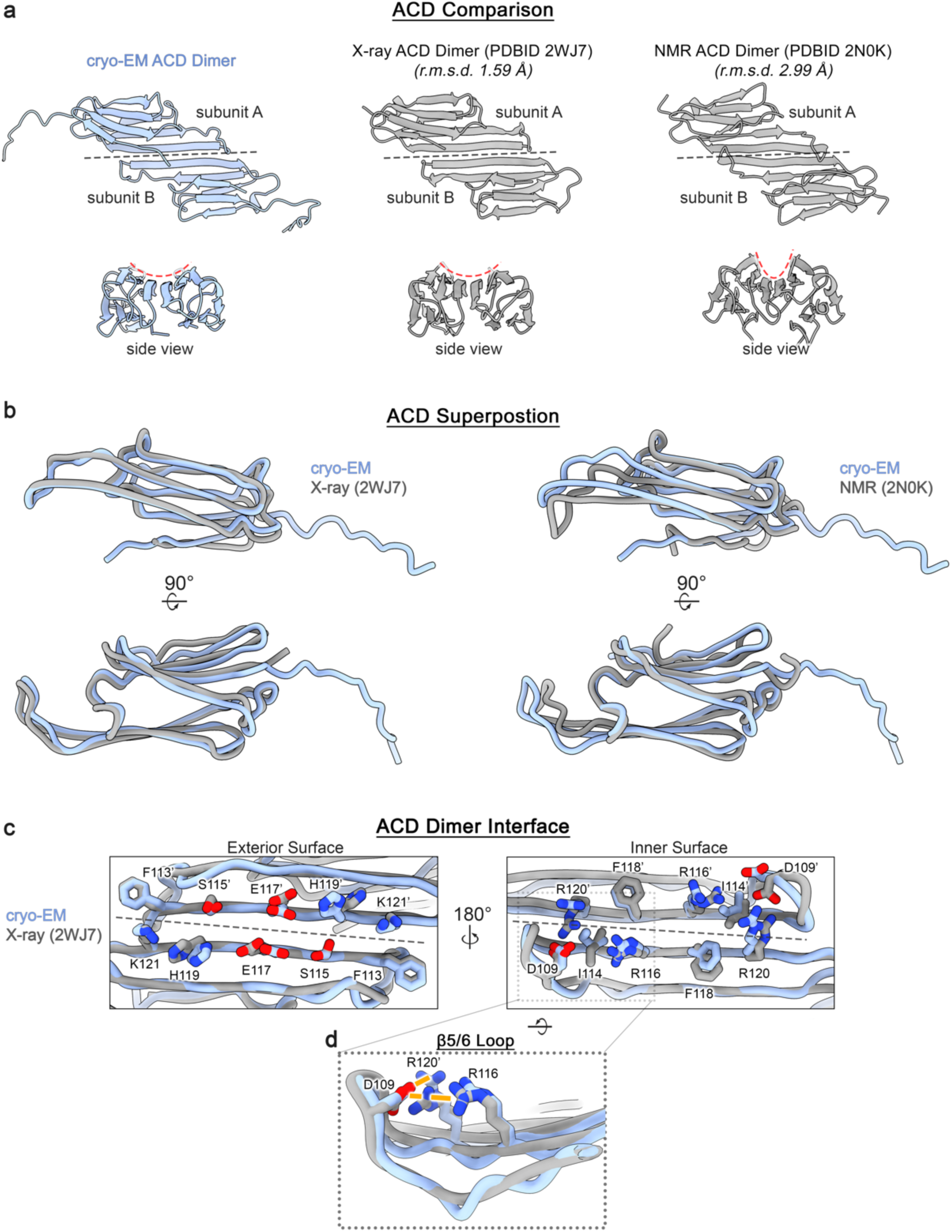
Comparison of structural features between αB-AXA and truncated αB-crystallin ACD models. **a.** Structural comparison of the αB-AXA ACD dimer (blue) to previously reported αB-crystallin ACD structures obtained X-ray crystallography (PDB: 2WJ7)^27^ (middle) and solution-state NMR (PDB: 2N0K)^31^ (right) in grey. Root-mean-square-deviation (r.m.s.d.) values displayed. Grey dotted line indicated dimer interface. Red dotted line indicates curvature of beta sheet formed by β4-6/7 across the ACD dimer. **b.** Structural superposition of the αB-AXA ACD (blue, subunit A) to the X-ray model (2WJ7, subunit A) (left) and solution-state NMR model (2N0K, subunit A) (right) in grey. **c.** Comparison of the ACD dimer interface as resolved in the αB-AXA cryo-EM ACD and the X-ray ACD dimer (PDB 2WJ7), showing a similar symmetric AP_II_ configuration. The dimer forming residues are displayed (stick representation, colored by hetero-atom) and numbered accordingly for the two protomers for the exterior surface (left) and interior surface (right), with respect to the helical fibril assembly of αB-AXA. Grey dotted line indicates the dimer interface. **d.** Zoom view of panel b (right), showing the β5/6 loop in the so-called upward conformation. This conformation appears to be stabilized by a conserved aspartate (D109) that forms electrostatic interactions (orange dotted lines) with R116 and R120’ from the neighboring protomer.

**Extended Data Figure 6.**
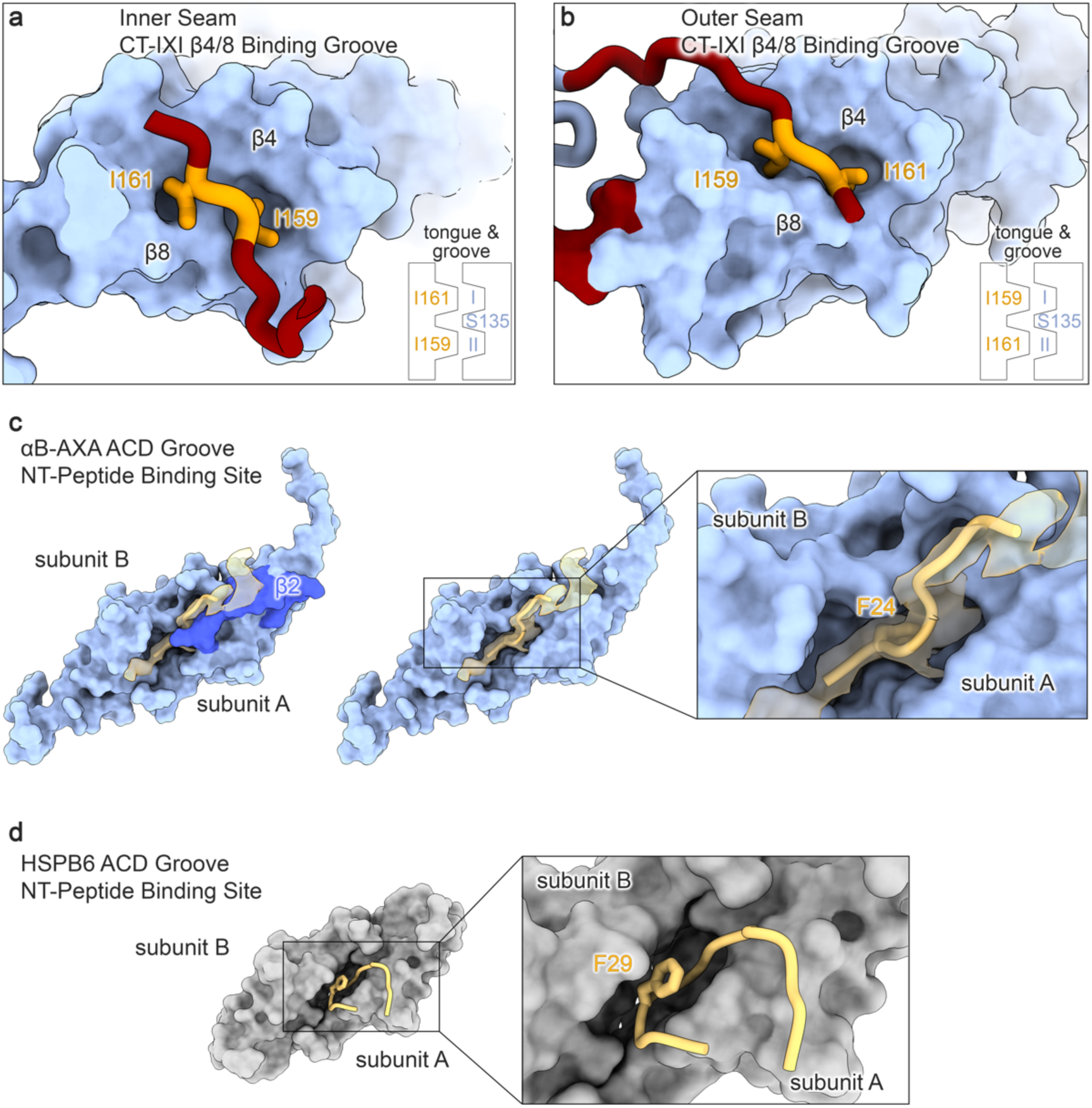
Comparison of αB-AXA CT- and NT-domain peptide binding sites on the α-crystallin domain. **a,b.** Different orientations of the palindromic CT-IXI motif bound to the hydrophobic β4/β8 groove of the ACD, where the inner seam interaction (panel a) adopts an anti-parallel (with respect to the β8 strand) orientation and the outer seam interaction involves a parallel orientation (panel b). The conserved isoleucines (I159 and I161) within the CT-IXI motif form a tongue and groove type fit, where hydrophobic residues in the ACD form two pockets (I and II) that are bifurcated by S135 in the ACD. These two pockets can accept the palindromic CT-IXI motif in either orientation. **c.** Putative interaction between the αB-AXA ACD dimer (blue) and the conserved hydrophobic region of the NTD (yellow). The β2 strand from a single protomer of the ACD (dark blue) lays across the NTD region (left), forming a lid-like feature that appears to stabilize the interaction with the ACD groove (right). The putative assignment this NTD region would place a conserved F24 residue into the hydrophobic groove of the ACD. **d.** A comparable interaction between the conserved NT region and the ACD of HSPB6 is observed in a previously reported crystallographic structure (PDB: 5LTW)^51^.

**Extended Data Figure 7.**
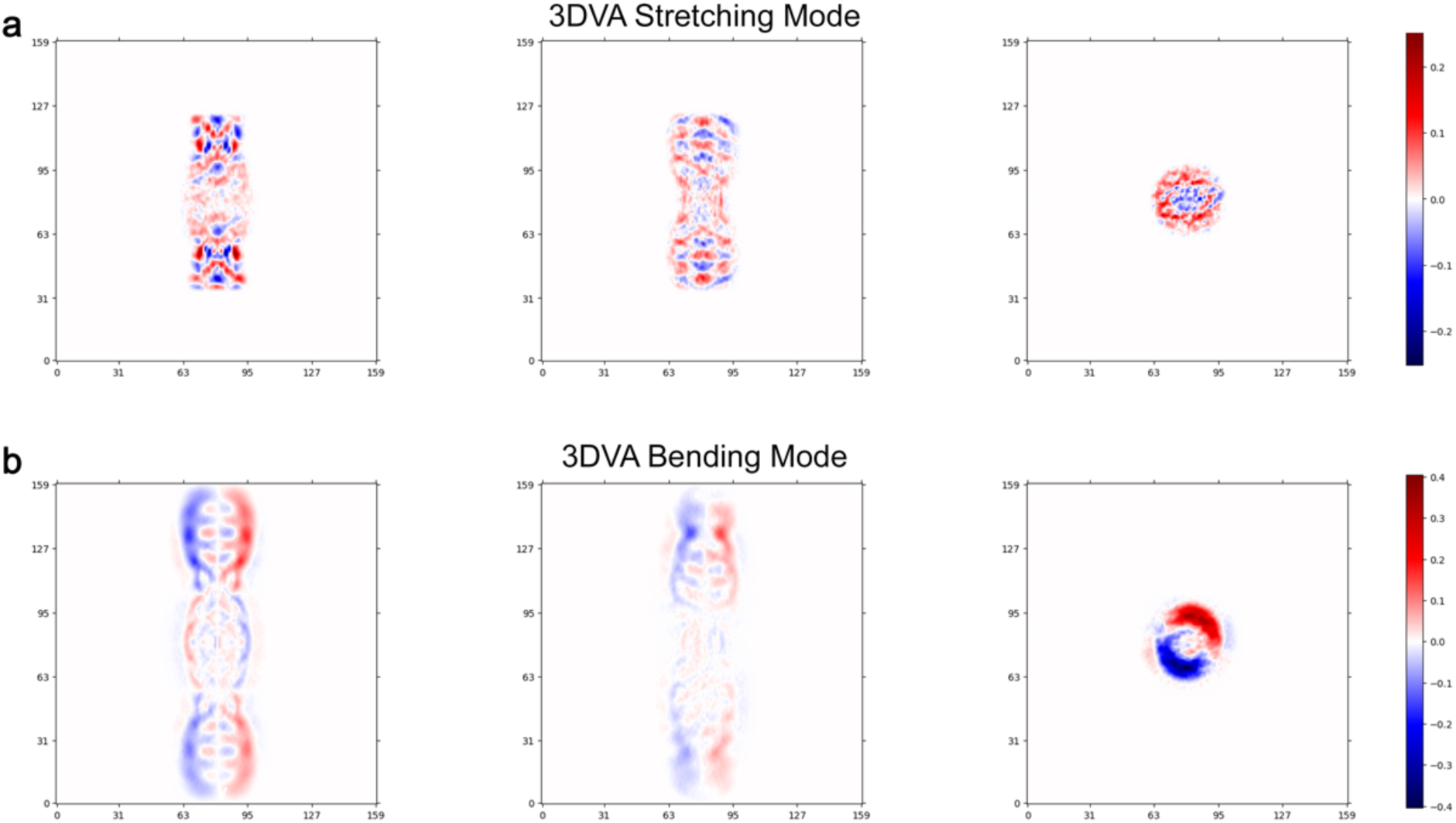
Eigenvectors obtained from 3D variability analysis (3DVA) in cryoSPARC. **a.** Heatmap illustrating three orthogonal slices of the identified eigenvector that describes a stretching mode. **b.** Heatmap illustrating three orthogonal views of the identified eigenvector that describes a bending mode.

## Supplemental Movie Legends

**Supplemental Movie 1. Cryo-EM 3D reconstruction and atomic model of αB-AXA fibril assembly.**

**Supplemental Movie 2. 3D variability analysis illustrating fibril bending and stretching modes.**

## REFERENCES

1. Horwitz, J. Alpha-crystallin can function as a molecular chaperone. Proc Natl Acad Sci U S A 89, 10449–53 (1992).

2. Jakob, U., Gaestel, M., Engel, K. & Buchner, J. Small heat shock proteins are molecular chaperones. J Biol Chem 268, 1517–20 (1993).

3. Reinle, K., Mogk, A. & Bukau, B. The Diverse Functions of Small Heat Shock Proteins in the Proteostasis Network. J Mol Biol 434, 167157 (2022).

4. Mogk, A., Bukau, B. & Kampinga, H.H. Cellular Handling of Protein Aggregates by Disaggregation Machines. Mol Cell 69, 214–226 (2018).

5. Kappe, G. et al. The human genome encodes 10 alpha-crystallin-related small heat shock proteins: HspB1-10. Cell Stress Chaperones 8, 53–61 (2003).

6. de Jong, W.W., Caspers, G.J. & Leunissen, J.A. Genealogy of the alpha-crystallin--small heat-shock protein superfamily. Int J Biol Macromol 22, 151–62 (1998).

7. Iwaki, T., Kume-Iwaki, A. & Goldman, J.E. Cellular distribution of alpha B-crystallin in non-lenticular tissues. J Histochem Cytochem 38, 31–9 (1990).

8. Slingsby, C. & Wistow, G.J. Functions of crystallins in and out of lens: roles in elongated and post-mitotic cells. Prog Biophys Mol Biol 115, 52–67 (2014).

9. Bakthisaran, R., Tangirala, R. & Rao Ch, M. Small heat shock proteins: Role in cellular functions and pathology. Biochim Biophys Acta 1854, 291–319 (2015).

10. Sarparanta, J., Jonson, P.H., Kawan, S. & Udd, B. Neuromuscular Diseases Due to Chaperone Mutations: A Review and Some New Results. Int J Mol Sci 21(2020).

11. Horwitz, J. Alpha crystallin: the quest for a homogeneous quaternary structure. Exp Eye Res 88, 190–4 (2009).

12. Haslbeck, M., Weinkauf, S. & Buchner, J. Small heat shock proteins: Simplicity meets complexity. J Biol Chem 294, 2121–2132 (2019).

13. Haley, D.A., Horwitz, J. & Stewart, P.L. The small heat-shock protein, alphaB-crystallin, has a variable quaternary structure. J Mol Biol 277, 27–35 (1998).

14. Aquilina, J.A., Benesch, J.L., Bateman, O.A., Slingsby, C. & Robinson, C.V. Polydispersity of a mammalian chaperone: mass spectrometry reveals the population of oligomers in alphaB-crystallin. Proc Natl Acad Sci U S A 100, 10611–6 (2003).

15. Inoue, R. et al. New insight into the dynamical system of alphaB-crystallin oligomers. Sci Rep 6, 29208 (2016).

16. Miller, A.P., O’Neill, S.E., Lampi, K.J. & Reichow, S.L. The alpha-crystallin Chaperones Undergo a Quasi-ordered Co-aggregation Process in Response to Saturating Client Interaction. J Mol Biol 436, 168499 (2024).

17. van den Oetelaar, P.J., van Someren, P.F., Thomson, J.A., Siezen, R.J. & Hoenders, H.J. A dynamic quaternary structure of bovine alpha-crystallin as indicated from intermolecular exchange of subunits. Biochemistry 29, 3488–93 (1990).

18. Bova, M.P., McHaourab, H.S., Han, Y. & Fung, B.K. Subunit exchange of small heat shock proteins. Analysis of oligomer formation of alphaA-crystallin and Hsp27 by fluorescence resonance energy transfer and site-directed truncations. J Biol Chem 275, 1035–42 (2000).

19. Baldwin, A.J. et al. Quaternary dynamics of alphaB-crystallin as a direct consequence of localised tertiary fluctuations in the C-terminus. J Mol Biol 413, 310–20 (2011).

20. Inoue, R. et al. Elucidation of the mechanism of subunit exchange in alphaB crystallin oligomers. Sci Rep 11, 2555 (2021).

21. Peschek, J. et al. Regulated structural transitions unleash the chaperone activity of alphaB-crystallin. Proc Natl Acad Sci U S A 110, E3780–9 (2013).

22. de Jong, W.W., Leunissen, J.A. & Voorter, C.E. Evolution of the alpha-crystallin/small heat-shock protein family. Mol Biol Evol 10, 103–26 (1993).

23. Kriehuber, T. et al. Independent evolution of the core domain and its flanking sequences in small heat shock proteins. FASEB J 24, 3633–42 (2010).

24. Jehle, S. et al. N-terminal domain of alphaB-crystallin provides a conformational switch for multimerization and structural heterogeneity. Proc Natl Acad Sci U S A 108, 6409–14 (2011).

25. Carver, J.A., Aquilina, J.A., Truscott, R.J. & Ralston, G.B. Identification by 1H NMR spectroscopy of flexible C-terminal extensions in bovine lens alpha-crystallin. FEBS Lett 311, 143–9 (1992).

26. Shi, J., Koteiche, H.A., McHaourab, H.S. & Stewart, P.L. Cryoelectron microscopy and EPR analysis of engineered symmetric and polydisperse Hsp16.5 assemblies reveals determinants of polydispersity and substrate binding. J Biol Chem 281, 40420–8 (2006).

27. Bagneris, C. et al. Crystal structures of alpha-crystallin domain dimers of alphaB-crystallin and Hsp20. J Mol Biol 392, 1242–52 (2009).

28. Laganowsky, A. et al. Crystal structures of truncated alphaA and alphaB crystallins reveal structural mechanisms of polydispersity important for eye lens function. Protein Sci 19, 1031–43 (2010).

29. Clark, A.R., Naylor, C.E., Bagneris, C., Keep, N.H. & Slingsby, C. Crystal structure of R120G disease mutant of human alphaB-crystallin domain dimer shows closure of a groove. J Mol Biol 408, 118–34 (2011).

30. Hochberg, G.K. et al. The structured core domain of alphaB-crystallin can prevent amyloid fibrillation and associated toxicity. Proc Natl Acad Sci U S A 111, E1562–70 (2014).

31. Rajagopal, P. et al. A conserved histidine modulates HSPB5 structure to trigger chaperone activity in response to stress-related acidosis. Elife 4(2015).

32. Treweek, T.M., Rekas, A., Walker, M.J. & Carver, J.A. A quantitative NMR spectroscopic examination of the flexibility of the C-terminal extensions of the molecular chaperones, alphaA- and alphaB-crystallin. Exp Eye Res 91, 691–9 (2010).

33. Pasta, S.Y., Raman, B., Ramakrishna, T. & Rao Ch, M. The IXI/V motif in the C-terminal extension of alpha-crystallins: alternative interactions and oligomeric assemblies. Mol Vis 10, 655–62 (2004).

34. Delbecq, S.P., Jehle, S. & Klevit, R. Binding determinants of the small heat shock protein, alphaB-crystallin: recognition of the ‘IxI’ motif. EMBO J 31, 4587–94 (2012).

35. Ghosh, J.G. & Clark, J.I. Insights into the domains required for dimerization and assembly of human alphaB crystallin. Protein Sci 14, 684–95 (2005).

36. Clouser, A.F. et al. Interplay of disordered and ordered regions of a human small heat shock protein yields an ensemble of ‘quasi-ordered’ states. Elife 8(2019).

37. Ghosh, J.G., Shenoy, A.K., Jr. & Clark, J.I. N- and C-Terminal motifs in human alphaB crystallin play an important role in the recognition, selection, and solubilization of substrates. Biochemistry 45, 13847–54 (2006).

38. Mainz, A. et al. The chaperone alphaB-crystallin uses different interfaces to capture an amorphous and an amyloid client. Nat Struct Mol Biol 22, 898–905 (2015).

39. Jehle, S. et al. Solid-state NMR and SAXS studies provide a structural basis for the activation of alphaB-crystallin oligomers. Nat Struct Mol Biol 17, 1037–42 (2010).

40. Peschek, J. et al. The eye lens chaperone alpha-crystallin forms defined globular assemblies. Proc Natl Acad Sci U S A 106, 13272–7 (2009).

41. Braun, N. et al. Multiple molecular architectures of the eye lens chaperone alphaB-crystallin elucidated by a triple hybrid approach. Proc Natl Acad Sci U S A 108, 20491–6 (2011).

42. Kaiser, C.J.O. et al. The structure and oxidation of the eye lens chaperone alphaA-crystallin. Nat Struct Mol Biol 26, 1141–1150 (2019).

43. Siezen, R.J., Bindels, J.G. & Hoenders, H.J. The quaternary structure of bovine alpha-crystallin. Effects of variation in alkaline pH, ionic strength, temperature and calcium ion concentration. Eur J Biochem 111, 435–44 (1980).

44. Siezen, R.J., Bindels, J.G. & Hoenders, H.J. The quaternary structure of bovine alpha-crystallin. Size and charge microheterogeneity: more than 1000 different hybrids? Eur J Biochem 91, 387–96 (1978).

45. Selivanova, O.M. & Galzitskaya, O.V. Structural and Functional Peculiarities of alpha-Crystallin. Biology (Basel) 9(2020).

46. Punjani, A., Rubinstein, J.L., Fleet, D.J. & Brubaker, M.A. cryoSPARC: algorithms for rapid unsupervised cryo-EM structure determination. Nat Methods 14, 290–296 (2017).

47. Kim, K.K., Kim, R. & Kim, S.H. Crystal structure of a small heat-shock protein. Nature 394, 595–9 (1998).

48. Haley, D.A., Bova, M.P., Huang, Q.L., McHaourab, H.S. & Stewart, P.L. Small heat-shock protein structures reveal a continuum from symmetric to variable assemblies. J Mol Biol 298, 261–72 (2000).

49. White, H.E. et al. Multiple distinct assemblies reveal conformational flexibility in the small heat shock protein Hsp26. Structure 14, 1197–204 (2006).

50. Shi, J. et al. Cryoelectron microscopy analysis of small heat shock protein 16.5 (Hsp16.5) complexes with T4 lysozyme reveals the structural basis of multimode binding. J Biol Chem 288, 4819–30 (2013).

51. Sluchanko, N.N. et al. Structural Basis for the Interaction of a Human Small Heat Shock Protein with the 14-3-3 Universal Signaling Regulator. Structure 25, 305–316 (2017).

52. Woods, C.N., Ulmer, L.D., Guttman, M., Bush, M.F. & Klevit, R.E. Disordered region encodes alpha-crystallin chaperone activity toward lens client gammaD-crystallin. Proc Natl Acad Sci U S A 120, e2213765120 (2023).

53. Punjani, A. & Fleet, D.J. 3D variability analysis: Resolving continuous flexibility and discrete heterogeneity from single particle cryo-EM. J Struct Biol 213, 107702 (2021).

54. Shi, X. et al. Small heat shock protein AgsA forms dynamic fibrils. FEBS Lett 585, 3396–402 (2011).

55. Vicart, P. et al. A missense mutation in the alphaB-crystallin chaperone gene causes a desmin-related myopathy. Nat Genet 20, 92–5 (1998).

56. Litt, M. et al. Autosomal dominant congenital cataract associated with a missense mutation in the human alpha crystallin gene CRYAA. Hum Mol Genet 7, 471–4 (1998).

57. Fichna, J.P. et al. A novel dominant D109A CRYAB mutation in a family with myofibrillar myopathy affects alphaB-crystallin structure. BBA Clin 7, 1–7 (2017).

58. Sacconi, S. et al. A novel CRYAB mutation resulting in multisystemic disease. Neuromuscul Disord 22, 66–72 (2012).

59. Bova, M.P. et al. Mutation R120G in alphaB-crystallin, which is linked to a desmin-related myopathy, results in an irregular structure and defective chaperone-like function. Proc Natl Acad Sci U S A 96, 6137–42 (1999).

60. Treweek, T.M. et al. R120G alphaB-crystallin promotes the unfolding of reduced alpha-lactalbumin and is inherently unstable. FEBS J 272, 711–24 (2005).

61. Hafizi, M. et al. Structural and functional studies of D109A human alphaB-crystallin contributing to the development of cataract and cardiomyopathy diseases. PLoS One 16, e0260306 (2021).

62. Ghahramani, M. et al. Structural and functional characterization of D109H and R69C mutant versions of human alphaB-crystallin: The biochemical pathomechanism underlying cataract and myopathy development. Int J Biol Macromol 146, 1142–1160 (2020).

63. Muhlhofer, M. et al. Phosphorylation activates the yeast small heat shock protein Hsp26 by weakening domain contacts in the oligomer ensemble. Nat Commun 12, 6697 (2021).

64. Biswas, S., Garg, P., Dutta, S. & Suguna, K. Multiple nanocages of a cyanophage small heat shock protein with icosahedral and octahedral symmetries. Sci Rep 11, 21023 (2021).

65. Hochberg, G.K. & Benesch, J.L. Dynamical structure of alphaB-crystallin. Prog Biophys Mol Biol 115, 11–20 (2014).

66. van Montfort, R.L., Basha, E., Friedrich, K.L., Slingsby, C. & Vierling, E. Crystal structure and assembly of a eukaryotic small heat shock protein. Nat Struct Biol 8, 1025–30 (2001).

67. Clark, A.R. et al. Terminal Regions Confer Plasticity to the Tetrameric Assembly of Human HspB2 and HspB3. J Mol Biol 430, 3297–3310 (2018).

68. Klevit, R.E. Peeking from behind the veil of enigma: emerging insights on small heat shock protein structure and function. Cell Stress Chaperones 25, 573–580 (2020).

69. Ghosh, J.G., Estrada, M.R. & Clark, J.I. Interactive domains for chaperone activity in the small heat shock protein, human alphaB crystallin. Biochemistry 44, 14854–69 (2005).

70. Bhattacharyya, J., Padmanabha Udupa, E.G., Wang, J. & Sharma, K.K. Mini-alphaB-crystallin: a functional element of alphaB-crystallin with chaperone-like activity. Biochemistry 45, 3069–76 (2006).

71. Pasta, S.Y., Raman, B., Ramakrishna, T. & Rao Ch, M. Role of the conserved SRLFDQFFG region of alpha-crystallin, a small heat shock protein. Effect on oligomeric size, subunit exchange, and chaperone-like activity. J Biol Chem 278, 51159–66 (2003).

72. Ahmad, M.F., Raman, B., Ramakrishna, T. & Rao Ch, M. Effect of phosphorylation on alpha B-crystallin: differences in stability, subunit exchange and chaperone activity of homo and mixed oligomers of alpha B-crystallin and its phosphorylation-mimicking mutant. J Mol Biol 375, 1040–51 (2008).

73. Schindelin, J., et al. Fiji: an open-source platform for biological-image analysis. Nat Methods 9, 676–82 (2012).

74. Tang, G. et al. EMAN2: an extensible image processing suite for electron microscopy. J Struct Biol 157, 38–46 (2007).

75. Scheres, S.H. RELION: implementation of a Bayesian approach to cryo-EM structure determination. J Struct Biol 180, 519–30 (2012).

76. Mastronarde, D.N. Automated electron microscope tomography using robust prediction of specimen movements. J Struct Biol 152, 36–51 (2005).

77. He, S. & Scheres, S.H.W. Helical reconstruction in RELION. J Struct Biol 198, 163–176 (2017).

78. Goddard, T.D. et al. UCSF ChimeraX: Meeting modern challenges in visualization and analysis. Protein Sci 27, 14–25 (2018).

79. Croll, T.I. ISOLDE: a physically realistic environment for model building into low-resolution electron-density maps. Acta Crystallogr D Struct Biol 74, 519–530 (2018).

80. Afonine, P.V. et al. Real-space refinement in PHENIX for cryo-EM and crystallography. Acta Crystallogr D Struct Biol 74, 531–544 (2018).

81. Emsley, P. & Cowtan, K. Coot: model-building tools for molecular graphics. Acta Crystallogr. D60, 2126–2132 (2004).

82. Williams, C.J. et al. MolProbity: More and better reference data for improved all-atom structure validation. Protein Sci 27, 293–315 (2018).

